# A single cell trajectory of human archetypal pluripotent stem cell differentiation to trophoblast stem cells reveals induction of endogenous BMP5/7 and GATA3 without transitioning through a naive state

**DOI:** 10.1101/2020.08.29.273425

**Authors:** Ethan Tietze, Andre Rocha Barbosa, Bruno Henrique Silva Araujo, Veronica Euclydes, Hyeon Jin Cho, Yong Kyu Lee, Arthur Feltrin, Bailey Spiegelberg, Alan Lorenzetti, Joyce van de Leemput, Pasquale Di Carlo, Tomoyo Sawada, Gianluca Ursini, Kynon J. Benjamin, Helena Brentani, Joel E. Kleinman, Thomas M. Hyde, Daniel R. Weinberger, Ronald McKay, Joo Heon Shin, Apua C.M. Paquola, Jennifer A. Erwin

**Affiliations:** Lieber Institute for Brain Development, Baltimore, Maryland, USA; Department of Neurology, Johns Hopkins School of Medicine, Baltimore, Maryland, USA; Inter-institutional Graduate Program on Bioinformatics, University of São Paulo, São Paulo, SP, Brazil; Center for Mathematics, Computation and Cognition, Federal University of ABC, Santo André, SP, Brazil; Department of Neuroscience, Johns Hopkins School of Medicine, Baltimore, Maryland, USA; Department of Psychiatry & Behavioral Sciences, Johns Hopkins University School of Medicine, Baltimore, MD, USA; Department of Genetic Medicine, Johns Hopkins University School of Medicine, Baltimore, MD, USA; Department of Psychiatry, University of Sao Paulo, Medical School, São Paulo, Brazil; Department of Basic Medical Science, Neuroscience, and Sense Organs – University of Bari Aldo Moro, Bari, Italy

**Author notes:** Center for Precision Disease Modeling and Division of Endocrinology, Diabetes and Nutrition, Department of Medicine, University of Maryland School of Medicine, 670 West Baltimore Street, Baltimore, MD, 21201, USA. Equal contributor (Ethan Tietze, Andre Rocha Barbosa, Bruno Henrique Silva Araujo, Veronica Euclydes).

**Keywords:** Placenta, Trophoblast, Trophectoderm, Primed iPSC, BMP

## Abstract

The human placenta is increasingly a focus of research related to early child development and the impact of maternal hyperimmune states. Primary human trophoblast stem cells (hTSC) and human pluripotent stem cells (hPSC) differentiated to hTSC can potentially model placental processes *in vitro*. Yet, it remains controversial how the differentiation of human pluripotent stem cells to trophectoderm relates to *in vivo* development and the factors required for this differentiation. Here, we demonstrate that the primed pluripotent state retains potency to generate trophoblast stem cells by activating EGF and WNT and inhibiting TGFb, HDAC and ROCK signaling without exogenous BMP4 (named TS). We map this specification by temporal single cell RNAseq compared to activating BMP4 or activating BMP4 and inhibiting WNT. TS conditions generate a stable proliferating cell type that is highly similar to six-week placental cytotrophoblasts with activation of endogenous retroviral genes and without amnion expression. Multiple primed iPSC and ES lines differentiate to iPS-derived-TSCs that can be passaged for at least 30 passages and differentiate to pure populations of multinucleated syncytiotrophoblasts and extravillous trophoblast cells. Our findings establish that primed iPS cell specification to hTSC with TS conditions involves induction of *TMSB4X*, *BMP5/7*, GATA3 and *TFAP2A* without transitioning through a naive state. Collectively, our results suggest that the primed state is on a continuum of potency and can differentiate to trophoblast stem cells via multiple paths.

**Significance Statement:** In the present study, we map the specification of primed induced pluripotent stem cells to trophoblast stem cells (TSC). Primed iPS-derived-TSC share transcriptional, morphological and functional characteristics with human *ex vivo* cytotrophoblasts including capacity of self-renewal and the ability to differentiate to pure extravillous and syncytiotrophoblasts. iPS-derived TSC display a uniquely active transcriptional network of human endogenous retroviruses similar to *in vivo* trophoblast. In addition, the fast conversion of primed iPSC to TSC allows for modeling placental diseases from large pluripotent stem cell cohorts which are traditionally banked at the primed state. Collectively, our results suggest that the primed state is on a continuum of potency which can differentiate to trophoblast stem cells via multiple paths.

## Introduction

Human models are essential to study human placenta development and function in health and disease. Comparative genome analysis demonstrates that genes involved in placenta and reproduction are amongst the most divergent between human and mouse [1]. For the human placenta, which is increasingly implicated as a risk factor for developmental disorders associated with a maternal hyperimmune state [2], the ability to model human trophectoderm *in vitro* is a powerful platform to identify disease-associated molecular targets, perform phenotypic screens and evaluate toxicity. *In vitro* and *in vivo*, trophoblast stem cells (TSC) are proliferative stem cell precursors that can differentiate into the trophectoderm cell types syncytiotrophoblasts and extravillous cytotrophoblasts [3].

Questions for how to appropriately model disorders in hTSC using hPSC cohorts exist [4]. Cohorts of iPSCs derived from patients can be used to model disease processes and genetically encoded traits. iPSCs cohorts and large iPSC banks are almost universally grown in primed conditions. Thus, the initial stage of most iPSC cohort studies will begin with cells in the primed state.

Conflicting results raise questions about the developmental potential and specification processes of hPSC to TSC [4]. Multiple culture conditions direct human stem cells to various developmental stages, including “naive” conditions which achieve a pre-implantation-like epiblast state [5, 6, 7] and archetypical “primed” conditions which achieve a post implantation-like epiblast state. It is widely accepted that the extraembryonic trophoblast lineage is specified before implantation. Therefore the differentiation of primed and naive pluripotent stem cells to trophectoderm does not normally occur *in vivo*. Questions remain about the heterogenous pluripotent states of archetypical primed PSC and the competency of these states to give rise to trophoblast cells [4]. Some studies suggest that BMP-treated primed hPSC resemble amnion and not trophoblast, while several more recent studies conclusively demonstrate that primed hPSCs can generate *bona fide* TSC with BMP4 treatment [8,9] or TGFb inhibition [10]. These differing reports may be reconciled by the fact that amnion specification in humans may be highly similar to trophectoderm [11] and that non-optimized culture conditions caused conflicting results [10].

Here, we sought to define the developmental trajectory of primed iPSC conversion to trophoblast stem cells and the involvement of exogenous BMP4 and WNT. The TS state can be reached via multiple paths. Four factor reprogramming of human fibroblasts cultured in either primed or naive conditions leads to upregulation of trophectoderm programs, and trophoblast stem cell lines can be derived from both naive and primed reprogramming intermediates [12]. Activn/Nodal, FGF, IGF and WNT signaling regulate pluripotency in human primed PSCs, and manipulating these pathways can induce primed PSC specification to trophectoderm and other embryonic lineages [13]. Bone Morphogenetic Protein 4 (BMP4) induces differentiation of human primed PSCs to trophoblast-like cells [14]. In the presence of BMP4, Inhibition of FGF and activin/nodal signaling [15, 16] or inhibition of WNT signaling generates more uniform trophectoderm [17] and trophoblast stem cells [8]. The differentiation in the presence of BMP4 follows a trajectory through a naive state [8]. Trophoblast can also be generated from human primed iPSC by activation of WNT and EGF while inhibiting Transforming Growth Factor Beta (TGFb), histone deacetylase (HDAC) and Rho-associated protein kinase (ROCK) [3], which is enhanced in the presence of BMP4 or by removing WNT activation [10].

In the present study, we find that activation of WNT and EGF while inhibiting TGFb, HDAC and ROCK (TS condition) in the absence of exogenous BMP4 specifies multiple primed pluripotent stem cell lines to TSC. We demonstrate that primed-derived-TSCs are capable of self-renewal for at least 30 passages and can differentiate into syncytiotrophoblast and extravillous trophoblasts. We used temporal single-cell sequencing analysis to elucidate the trajectory of this specification. Trophoblast cells generated by TS condition are transcriptionally highly similar to *ex vivo* cytotrophoblasts and uniquely upregulate endogenous retroviral genes. The specification employs established trophectoderm programs including *TMSB4X*, *YAP*, *GATA3, CDX2* and *TFAP2A* and interestingly induces endogenous *BMP5* and *BMP7* expression, indicating a role for endogenous BMP signaling even in the absence of exogenous BMP4 treatment.

## Results

### Derivation of proliferative trophoblast-like stem cells (TSC) from primed human embryonic stem cells and primed induced pluripotent stem cells without BMP4

We asked if primed ESC and iPSC can differentiate to TSC without exogenous BMP4 in the conditions used to generate trophoblast stem cells from blastocysts and human villus cytotrophoblasts [3], here referred to as TS which activates EGF and WNT and inhibits TGFb, HDAC and ROCK signaling. While previously described primed iPSC differentiations to trophectoderm have added BMP4, evidence has shown that iPSC can differentiate to trophectoderm in the absence of exogenous BMP4 [18]. 24 hours after clump passage of the human embryonic stem cell line H1 [19] in primed feeder-free stem cell conditions using StemFlex, ES colonies demonstrated tight packing with defined edges characteristic of primed cells (Figure 1a and b). Once self organized colonies were established 24 hours post passage, cells were switched directly to TS condition media. By brightfield imaging over the subsequent 6 days, we observed that cells proliferated rapidly and adopted a flatter appearance with some cells adopting a cobblestone appearance (Figure 1b-d). After subsequent passaging, two morphologically distinct populations emerged. Circular colonies with an epithelial-like appearance were surrounded by phase bright fibroblastic cells (Figure 1e,f). The inner epithelial-like cells continued to proliferate and appeared morphologically similar to TSC derived from blastocysts and human villus cytotrophoblasts (Figure 1g,j,l and n). Immunofluorescence revealed that subpopulations of the inner cells expressed TP63 and/or KRT7 and minimally expressed VIM while surrounding fibroblastic cells strongly expressed VIM (Figure 1h). After several passages, the epithelial-like cells preferentially expand and reach purity between passage 5-7. To confirm the ability to proliferate as TSC-like cells, iPS-derived trophoblast stem cell lTSC lines were passed for up to 32 passages and maintained KRT7 and TP63 expression (Figure 1i). These TSC widely express both TP63 and KRT7 (Figure 1j-o, Figure S1) without expressing the pluripotency genes SOX2 and NANOG (Figure S1k-n). In addition, the majority of cells highly express ITGA6 (89%), a common cell surface marker that is present in cytotrophoblast and human trophoblast stem cells [3] (Figure 1p).

**Figure 1:**
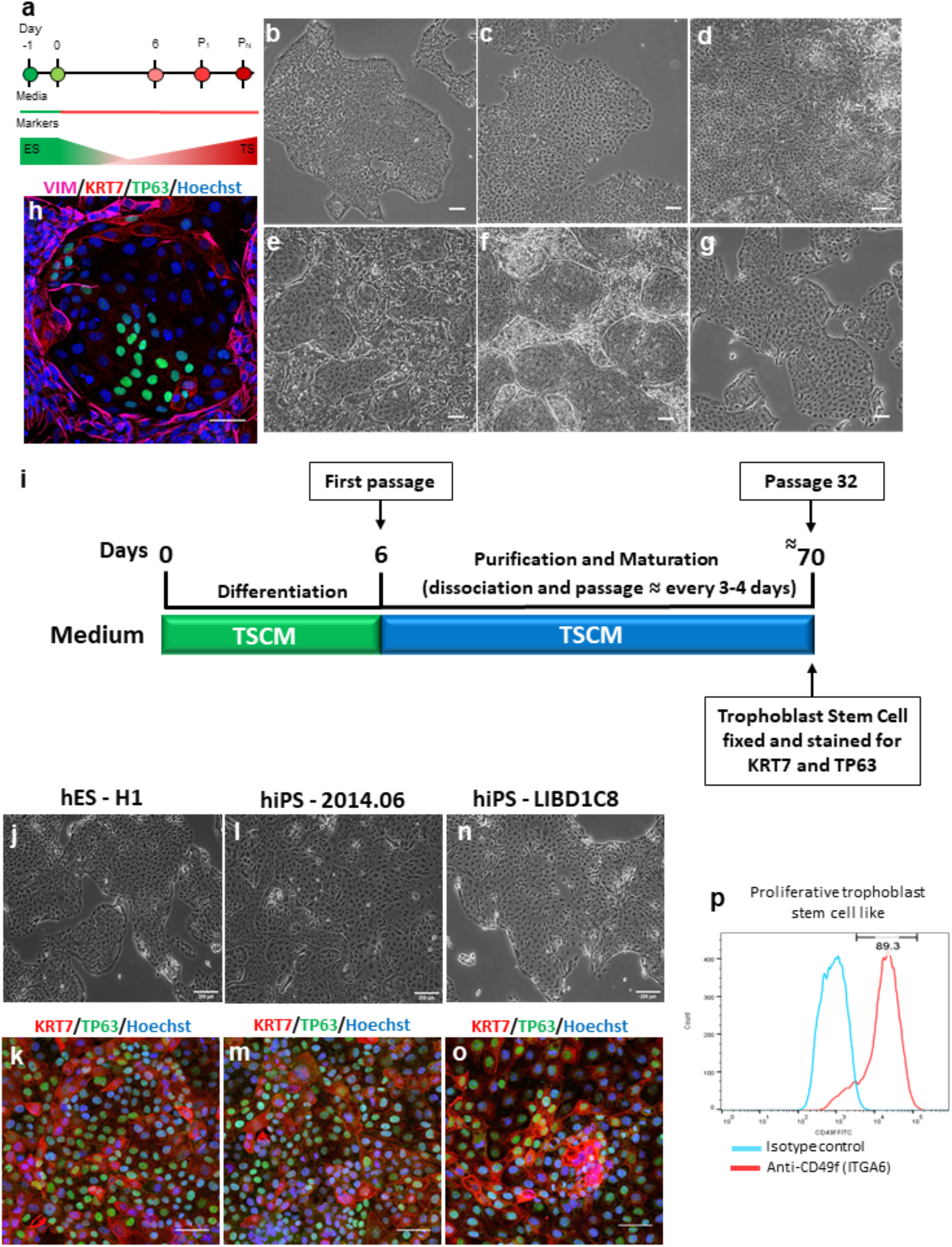
Derivation of Trophoblast Stem Cell Like (TS) Cells From Human Pluripotent Stem Cells. **(a).** Schematic of protocol used to derive TS cells from human pluripotent stem cells by activating wingless/Integrated (WNT) and epidermal growth factor (EGF) and inhibiting Transforming Growth Factor Beta (TGFb) histone deacetylase (HDAC) and Rho-associated protein kinase (ROCK). **(b-d).** Representative phase contrast images of differentiating TS cells (ES line H1). Images of differentiating cells on days 2 (b), 4 (c), and 6 (d) after addition of differentiation (TS) media. **(e,f).** Images of TS cell morphology 24 (e) and 72 (f) hours following passage (P1) on day 6. **(g).** Mature TS lines generated from ES line H1 after 10 passages. **(h).** Immunofluorescence for VIM (magenta), KRT7 (red), TP63 (green), and Hoechst 33342 (blue) in TS cells after 1 passage (P_0_)(iPSC line LIBD7c6). **(i).** Schematic representation of the differentiation protocol of iPSC to TSC. TSCM = Trophoblast Stem Cell Medium. **(j,l,n)**. Representative phase contrast images of TS cells H1 (j), 2014.06 (l) and LIBD1C8 (n). **(k,m,o).** Confocal images of KRT7 (red), TP63 (green), and Hoechst 33342 (blue) for TS cells generated from passage 11 H1 (k) and 2014.06 (m) and passage 32 LIBD1C8 (o). **(p).** Flow cytometry histograms of ITGA6 expression in mature TSC derived from H1 line. All phase contrast images scale bars indicate 100 µm. All immunofluorescence images scale bars indicate 50 µm.

Pluripotent stem cell lines vary in their propensity to differentiate to different cell types [20]. Therefore, we asked if TS condition could specify a variety of primed human stem cell lines from different sources. We confirmed TSC specification by VIM, KRT7 and TP63 immunofluorescence on iPSC lines reprogrammed from dermal fibroblasts by sendai, named 2014.06 (Supplemental Figure 2 and Figure S1k-r), and two postmortem dura fibroblast-derived iPS cell lines reprogrammed by episome, named LIBD1c8 and LIBD7c6 [21] (Figure 1n and o and S1a-j).

### Differentiation of Trophoblast Cells into multinucleated syncytiotrophoblasts and extravillous trophoblast cells

To investigate the differentiation potential of primed-derived-TSC, we evaluated the ability of primed-derived-TSC to differentiate to mature trophectoderm cell types. Proliferative trophoblast stem cells are bipotential stem cells with the capacity to differentiate into syncytiotrophoblast (STB) and extravillous trophoblasts (EVT) [3]. EVT differentiation of the primed-derived-TSC H1 line was induced with the TGFP-B inhibitor A83, NRG1 and Matrigel, as previously described for TSC (Figure 2a). After 9 days in EVT differentiation, primed-derived-TSC acquired EVT morphology (Figure 2b) and 88% of cells were positive for the EVT specific marker HLA-G by immunofluorescence staining (Figure 2c,d), thereby resembling previously described derived human TSC [3].

**Figure 2:**
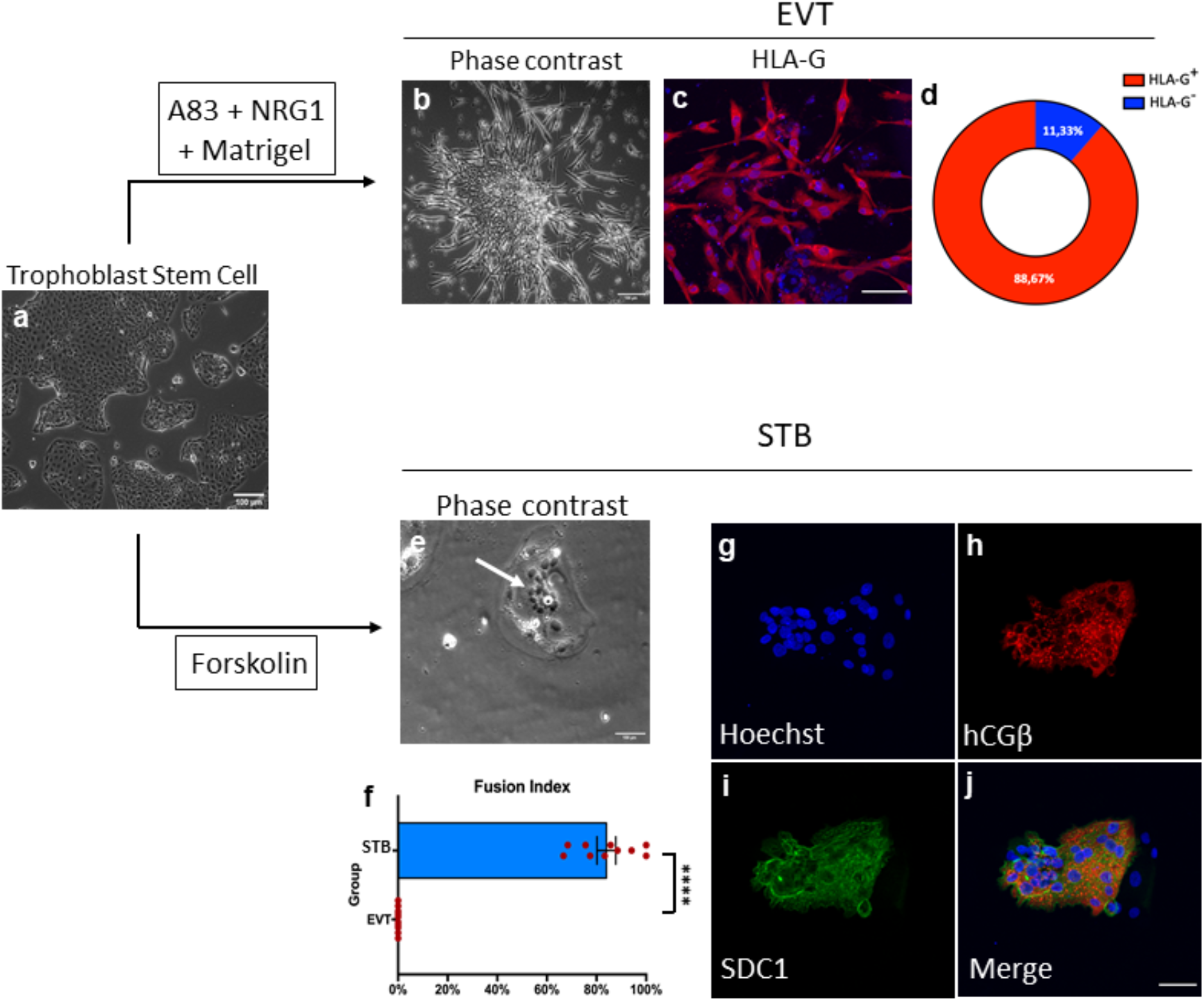
iPS-derived TS Cells Differentiate Into Syncytiotrophoblast, Extravillous Cytotrophoblasts and Villi-like Structures. **(a).** Representative phase contrast images of differentiating TS cells. Scale bars indicate 200 µm **(b).** Representative image of EVT derived cells. Scale bars indicate 100 µm. **(c).** Immunofluorescence image of EVT derived cell from mature TSC line H1 stained for Hoechst 33342 (blue) and HLA-G (red). Scale bars indicate 100 µm. **(d).** Percentage of HLA-G positive cells after 6 days of differentiation. Scale bars indicate 100 µm. **(e).** Representative image of derived STB. Arrow indicates the multinucleated ST characteristic. Scale bars indicate 100 µm. **(f).** Fusion efficiency of STB. H1 ESC - STs were analyzed. Mean + SEM. ∗∗∗∗p ≤ 0.0001 by Wilcoxon-Mann-Whitney test.

To confirm differentiation to syncytiotrophoblast (STB), we induced the Primed-derived-TSC differentiation to STB with Forskolin for 3 days as previously described [3]. STB is a terminally differentiated multinucleated epithelial layer that infiltrates the maternal endometrium. The multinucleated cell forms from multiple cytotrophoblast cell fusions. Cells demonstrate STB morphology of multinucleation (Figure 2e) and the fusion index revealed that more than 80% of the cells were multinucleated (Figure 2f), similar to STB derived trophoblast cells from human villous cytotrophoblast and from the blastocyst outgrowth [3] and naive TSC [22]. Primed-TSC-derived STB expressed STB specific proteins hCGβ and syndecan-1 (SDC1) (Figure 2g-j), which were not detected in iPS cells (Supplemental Figure 1i). In summary, primed-derived-TSC are bipotential stem cells capable of efficient differentiation into both HLA-G positive villus cytotrophoblasts and hCGβ and SDC1 positive syncytiotrophoblasts.

### Single cell RNA sequencing reveals unique transcriptional programs for primed iPSC TSC specification in the absence of BMP4

To define the molecular events involved in the specification of primed PSC to trophectoderm, we performed temporal single cell RNAseq analysis of human iPSC specified the TSC using the previously described TS condition which activate EGF and WNT and inhibit TGFb, HDAC and ROCK signaling. Due to previous reports of BMP4 and Wnt involvement, we compared this specification to BMP4 treatment [14] and BMP4 and inhibition of WNT with the small molecule inhibitor IWP2 [17]. 24 hours after clump passaging with EDTA in StemFlex media, the media was changed to differentiation conditions. Single cells were sequenced at iPS cell stage before differentiation (day 0), at day 4 and day 6 after BMP4 treatment; on days 2, 4 and 6 for BMP4 + IWP2 treatment, and on days 2, 4, 6 and 8 for TS condition, resulting in 10 single-cell transcriptomes (Figure 3a). In TS condition, the day 8 is a timepoint two days after the first passage annotated as passage 1 (p1). In the BMP4 and BMP4 + IWP2 conditions, no proliferative cells survived passaging, therefore the p1 time point was not sequenced. Importantly we sought to compare the effects of altering BMP4 and WNT but do not aim to compare previously described hPSC specification methods; therefore, all differentiations began with iPS in stem flex media after clump passaging that gave rise to hTSC from multiple iPS lines (see Figure 2) but is a different iPS media from previous BMP4 protocols (see discussion). Using the highly parallel droplet based single cell sequencing method Drop-Seq [23], 9,821 high-quality cells were obtained after removing cells with less than 1000 genes detected and more than 20% of mitochondrial mapping. The average ratio of mapping reads for mitochondrial genes in all cells were 3.9%, indicating good viability (Supplemental Figure 3a). The total number of genes detected ranged between 21804 for BMP4 + IWP2 day 4 to 27134 for TS day 4 (Supplemental Figure 3b) time point with between 563 to 1572 high quality cells per time point (Supplemental Figure 3c).

**Figure 3:**
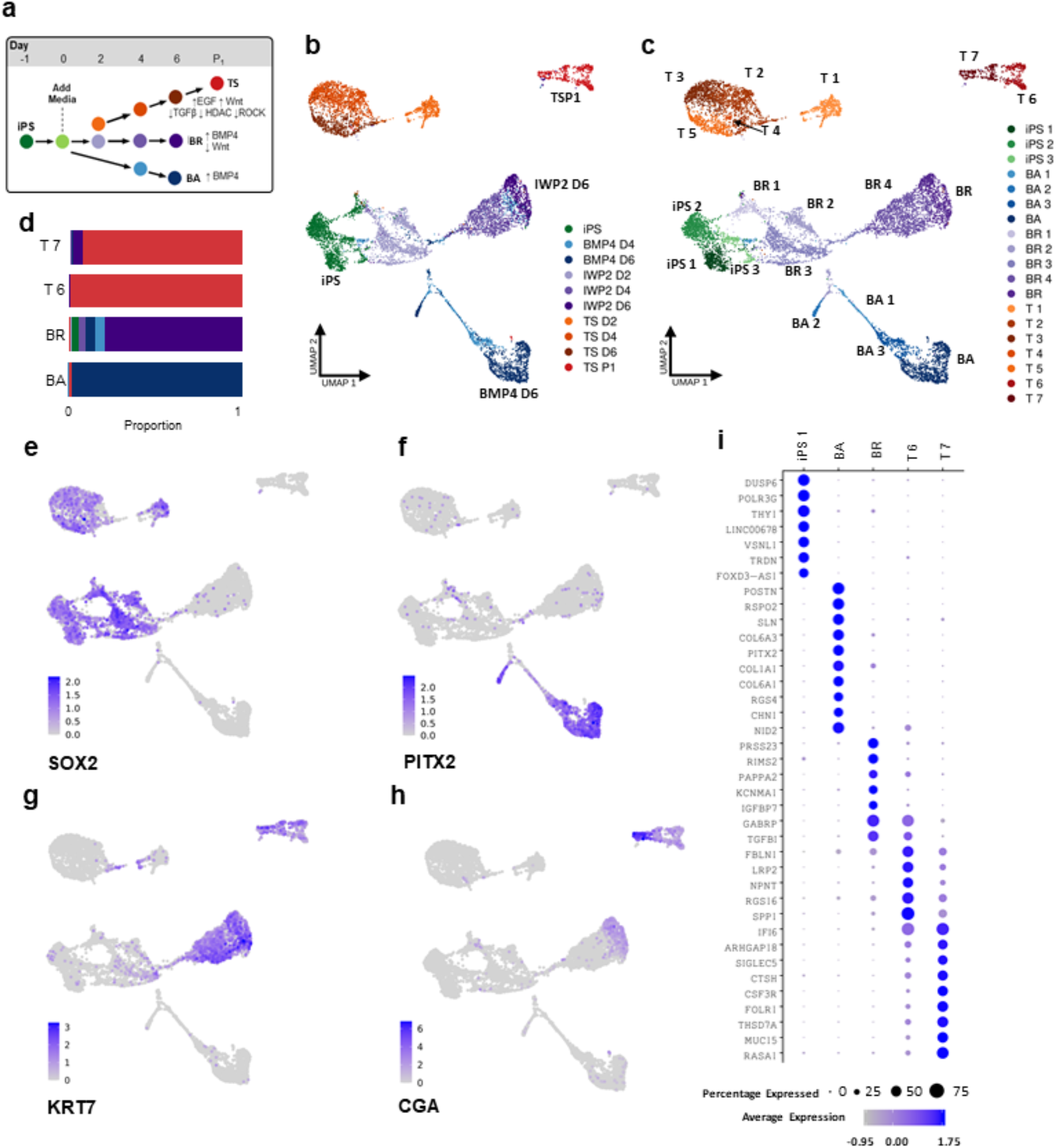
Temporal Single Cell RNA Sequencing Of iPSC To Trophoblast Differentiation. **(a).** Experimental schematic of trophoblast differentiation experiment. hiPS cells were plated and allowed to adhere to culture dishes for 24 hours before addition of differentiation media. Cells were collected for single-cell RNA sequencing at indicated time points. **(b,c).** Dimensionality reduction for the most variable genes across all cells with Principal Component Analysis (PCA) and Uniform Manifold Approximation and Projection (UMAP) embedding of 9,821 single cell transcriptomes from three distinct media treatments with groupings based on (b) sample identity or (c) gene expression clusters calculated by k nearest neighbors using the Euclidean distance of the 30 first PCs which identifies 19 clusters. Cells from the BMP4 condition almost exclusively formed four clusters termed BA (BMP4 alone cell indicated in blue). Cells from the BMP4+ IWP-2 condition were found predominantly in a group of five heterogeneous clusters termed BI (BMP responsive indicated in purple). iPSC indicated in green and TS clusters are indicated in orange. **(d).** Plot of the proportion of cells from each differentiation condition which are present in the final k-means clusters TS, T6, BI5 and BA4. Color represents the differentiation condition scale in 3B BMP4+IWP2: purple, BMP4: blue, iPSC: green, TS: orange. Cell proportions for all clusters can be found in supplemental table S1. **(e-h).** Single cell visualization of gene expression of selected markers of hiPS cells (e), placental stromal cells (f), and trophoblasts (g,h) on UMAP projections. Grey dots indicate cells with no expression and blue scale indicates normalized expression. **(i).** Dot plot displaying differentially upregulated marker genes for the most mature cell clusters from each treatment BA (BMP4 Alone), BI (BMP4+IWP2), and TS 6, TS (TS media) compared to the average expression of all other clusters. Size of dot indicates the percent of cells with expression and intensity of the blue indicates the average expression level.

We performed dimensionality reduction for the most variable genes across all cells with Principal Component Analysis (PCA) and Uniform Manifold Approximation and Projection (UMAP) embedding. Generally, single cells are separated by differentiation stage and condition, with UMAP dimension 2 generally capturing the differentiation condition (Figure 3b). To identify transcriptionally similar groups of cells (hereafter referred to as clusters), we performed a graph-based clustering analysis of k nearest neighbors using the euclidean distance of the 30 first PCs in Seurat [24]. We identified 19 clusters, and clusters contained mostly a single time-point and media condition (Figure 3b-d and S3d and Table S1). iPSC clusters 1, 2 and 3 mostly contained iPS cells with slight contributions from the first days of all three differentiation conditions. Cells from the BMP4 condition almost exclusively formed four clusters (termed <u>B</u>MP <u>A</u>lone) BA1, BA2, BA3 and BA4. BA1 and BA3 were predominantly composed of cells from day 4 of the differentiation while BA2 and BA4 were predominantly from day 6 cells. Cells from the BMP4+ IWP2 condition were found predominantly in a group of five heterogeneous clusters (termed <u>B</u>MP IWP2) BI1, BI2, BI3, BI4 and BI5. BI1 cluster was composed of roughly equal parts iPSCs and BMP4 + IWP2 day 2 treated cells. BMP4 + IWP2 cells made up the majority of the cells in the other BIs clusters alongside a smaller population of BMP4 condition. BI2 and BI3 clusters were predominantly derived from BMP4 + IWP2 day 2 cells, while day 4 contributed to BI4 cluster, and day 6 contributed to BI5. In addition, cluster BI5 contained a small contribution of cells originating from the final differentiation day of the TS condition termed T7 cluster, demonstrating capacity to differentiate to TSCs. Cells from the TS condition almost exclusively populated 7 clusters termed T1, T2, T3, T4, T5, T6 and T7 clusters. T1 cluster was exclusively composed of day 2 cells and T 2-5 clusters were composed of a mixture of cells from days 2-6. The final day of TS condition differentiation from cells 24 hours after passaging separated into two clusters named T6 and T7, which confirms the brightfield observations of 2 cell populations. Interestingly, 14 cells from BMP4+IWP2 day6 and 4 cells from the BMP4 day6 contribute to the T7 cluster (Figure 3d, Table S1).

To understand transcriptional changes during the differentiation, we asked which cells express canonical pluripotent, trophectoderm and mesoderm specific genes. The stem cell transcription factor *SOX2* is highest expressed in the iPSC stage and is absent from the most differentiated clusters (BA4, BI5 and T7) in all conditions (Figure 3e). *SOX2* expression is maintained at a higher level in the initial days of the TS condition compared to BMP4 + IWP2 and BMP4 treatment and absent from the most differentiated trophoblast stem cells. The Paired-like homeodomain transcription factor 2 PITX2 is highly expressed in the later stages of BMP4 differentiation (BA4 clusters), suggesting that BMP4 conditions generate cells with more mesenchyme phenotype (Figure 3f). In single cell RNAseq of the human placenta, *PITX2* was identified as a marker gene for a stromal fetal communicating cell type F1 by Vento-Tormo [25, 26]. Keratin 7, *KRT7*, and human chorionic gonadotropin subunit alpha, *CGA*, are expressed in both stem cell derived cytotrophoblasts and *in vivo* placenta [27]. Indicating specification to trophectoderm transcriptional programs, we find both *KRT7* and *CGA* are expressed in the later stages of the BMP4 + IWP2 condition (BI4 and BI5 clusters) and TS condition (T6 and T7 clusters) (Figure 3g-h and Figure S4d and e). Overall, both TS condition and BMP4 + IWP2 treated cells demonstrated an increased expression for most trophoblast markers including *GATA3* and *TFAP2C* (Figure S4a) while the BMP4 condition demonstrated a more mesoderm associated expression of *GATA4*, *TBXT* and *PDGFRA* (Figure S4b). In mouse, the Caudal-type homeobox 2 (CDX2) is involved in the segregation of the inner cell mass and trophectoderm lineages at the blastocyst stage by repressing OCT4/POU5F1 and NANOG in the trophectoderm. Overexpression of CDX2 in mouse embryonic stem cells causes differentiation to trophoblast stem cells [28, 29]. In humans, *CDX2* expression is initiated after blastocyst formation and has variable expression patterns in trophectoderm [30]. Similar to human embryo and TSC derived from human blastocyst outgrowth [3], *CDX2* is transiently expressed during the intermediate stages of specification (day2-4) for all conditions and is not expressed in the final TS condition state (Figure S4a).

Next we asked which genes are preferentially expressed in each cluster by non-parametric Wilcoxon rank sum test (adj.p-value < 0.05; logFC > 0.25), by comparing gene expression of cells in a cluster compared to all other cells (Figure 3i, Table S2). *DUSP6* and *THY1* are primed pluripotent stem cell state genes and are among the top marker genes identified in the iPS cells. The BA cluster genes include WNT signaling pathway *RSPO2* a WNT agonist [31] and *PITX2,* which interacts with WNT signaling and regulates collagen expression [32]. BI5 cluster genes include *PRSS23* a serine protease involved in Snail-dependent Epithelial to Mesenchymal Transition [33] and Insulin-like growth factor signaling *IGFBP7* [34] and *PAPPA2* [35]. Markers of the T7 cluster include EGFR signaling *IFI6*, and *MUC15* [36] and the YAP downstream effector *ARHGAP18* [37], recently implicated in the specification hiPSC to bipotent trophoblast stem cells [38].

In sum, activation of WNT and EGF and inhibition of TGFb, HDAC and ROCK of primed iPSC induces transcriptional programs that closely related trophoblast-like cells. The differentially expressed genes for all pairwise cluster comparisons are provided in Table S3.

### Trophectoderm expression signatures are enriched in iPS-derived trophoblast cells

To compare the iPS-derived cell types to peri-implantation human embryos, we calculated the scaled transcriptional similarity of our scRNAseq data to previously annotated gene signatures for epiblast, hypoblast and trophoblast from cultured human embryos [39] (Figure 4a-d). Two iPSC clusters exhibited the highest similarity to epiblast marker gene expression, with average score of 0.1 (Figure 4b). Interestingly, a population of cells from the BA2 cluster were most similar to hypoblast tissue profiles with a similarity score of 0.3 (Figure 4c). The most mature trophoblast T7 cluster was specific for trophoblast tissue expression with a transcriptional similarity of 0.6 (Figure 4d). The post-passaging TS condition cluster T6 and T7 are enriched in trophoblast gene expression. The T7 cluster contains the highest trophoblast-genes expression, as evaluated by pairwise comparison of the average trophoblast gene expression for all clusters (Wilcoxon Rank Sum Test with Holm-middle combined p value = 3.5e-41). Further confirming the trophoblast transcriptional similarity by gene set enrichment analysis, cluster T7 exhibited the highest enrichment for expression of trophoblast-signature genes (GSEA adjusted p value = 0.001, Figure S3a). In summary, the single cell expression profile of cells treated with TS condition gain a trophoblast identity 8 days after specification and passaging.

**Figure 4:**
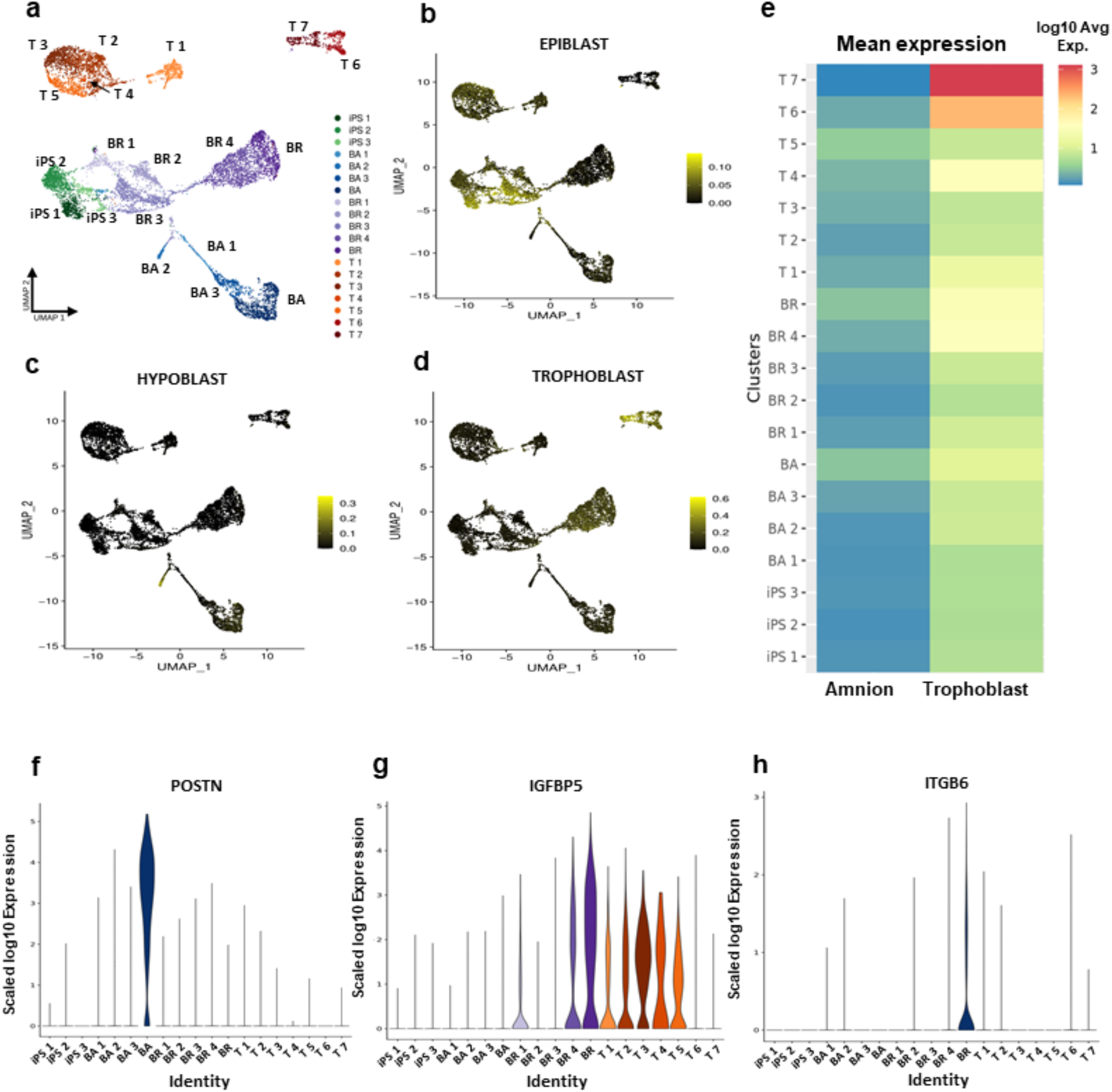
iPS-derived trophoblast cells specified without BMP4 are enriched for trophectoderm expression and are depleted for amnion expression. **(a).** UMAP of single cell transcriptomes from three distinct conditions with cluster labels. **(b,c,d).** Expression similarity score for single cells (a) based on gene signatures from Epiblast, Hypoblast and Trophoblast of human embryo [39]. (b) Two iPSC day 0 clusters exhibited the highest similarity to epiblast (average score of 0.1). (c) SC day 4 treatment was the most similar to hypoblast (similarity score of 0.3). (d) The T7 cluster was the most specific for trophoblast tissue (similarity score of 0.6). **(e).** Mean expression of previously identified amnion [42] and Trophoblast gene lists [39] in each cluster. The T7 cluster contains the higher average trophoblast-gene expression (Wilcoxon Rank Sum Test with Holm-middle combined p value = 3.5e-41) and also the lowest average expression for the amnion genes (Wilcoxon Rank Sum Test with Holm-middle combined p value = 4.8e-06), compared to all other cell clusters. **(f,g,h).** Violin plot indicating the expression distribution of *POSTN, IGFBP5 and ITGB6,* amnion markers, in all clusters. (f) *The POSTN* expression is significantly upregulated in the BA cluster (adjusted p value < 2.225074e-308). (g) The *IGFBP5* is significantly upregulated in BI5 (adjusted p value = 1.909090e-147) and T3 (adjusted p value = 4.301357e-148) clusters. (h) *ITGB6* expression is significantly upregulated in the BI5 cluster (adjusted p value = 8.890e-286).

Next, we compared the specification map with *in vivo* amnion. Recent studies suggest that primed pluripotent stem cells patterned with BMP4 give rise to amnion-like cells [22, 40, 41]. Importantly, genes expressed predominantly in human amnion were not enriched in cluster T7 containing mostly differentiated TS treated cells (GSEA adjusted p value = 1, Figure S5b,c), instead cluster T7 is most enriched for trophoblast genes (Figure 4e). We evaluated the expression of 198 genes predominantly expressed in human amnion [42] and previously reported to be upregulated in primed BMP4 condition [22]. The T7 cluster have the lowest average expression for the 198 amnion genes with 0.068 log normalized average expression compared to all other cell clusters (Wilcoxon Rank Sum Test with Holm-middle combined p value = 4.8e-06), compared to 0.47 for the BA cluster BMP4 treated cells, 0.46 for the BMP4 +IWP2 treated cells, and 0.16, 0.13 and 0.16 for the iPSC clusters (Figure 4e, S6, Table S4, S5 and S6). Instead, amnion genes are expressed in clusters from the earlier stages of TS condition specification and BMP4 treatment. For example, *POSTN* gene is significantly upregulated in cluster BA4 (adjusted p value < 2.225074e-308), which predominantly contains d6 BMP4 alone treated cells (Figure 4f). Moreover, the amnion gene IGFBP5 is significantly upregulated four and six days after differentiation in both TS and BPM4+IWP2 conditions (BI 4 adjusted p value = 4.389994e-95 and BI5 adjusted p value = 1.909090e-147, mostly containing day 4 and 6 BMP4 + IWP2 cells; and T3 cluster adjusted p value = 4.301357e-148; mostly containing day 4 TS treatment) (Figure 4g). Cinkornpumin et al report high expression of the amnion gene *ITGB6* in trophoblast cells derived from primed hPSC [40]. *ITGB6* is nearly absent from all TS condition single cells while the BMP4 + IWP2 D6 (Wilcoxon Rank Sum test adjusted p value = 9.332e-264) sample and BI5 cluster (Wilcoxon Rank Sum test adjusted p value = 8.890e-286) show significant higher upregulation (Figure 4h). In summary, 8 days after specification and passaging in TS condition, primed iPS cells are specified to trophoblast without amnion expression.

### iPS-derived trophoblast cells resemble first trimester placental cytotrophoblasts

We next asked how similar are the iPS-derived placental cells to *ex vivo* cells found in human first trimester placenta. To characterize the expression profiles of the 19 clusters, we analyzed the overlap with cell type-specific genes identified by two published single-cell studies of early human placenta [25, 26]. We performed gene set enrichment analyses using Cell-Specific Expression Analysis (CSEA) [43]. We identified cell type specific genes that were preferentially expressed in each cell type/cluster of the 19 clusters from iPS-derived cells by the specificity index probability (pSI) statistic at thresholds of p< 0.05. Next, we tested whether cell-type specific genes previously identified by single cell studies of human placenta are over-represented in the cell-type specific genes from our iPSC specification by hypergeometric test and applied the Bonferroni correction for multiple comparisons, considering all the tested gene lists [α = 0.05/ (19× (38+14)) = 5.1×10^-5^]. Vento-Tormo identified a set of 38 cell types from maternal (decidua and blood) and fetal tissues (placenta) of first trimester placenta and reported 30 cell-type specific genes per cell type [26]. We found that the transcriptional profile from all iPSC clusters and clusters from d2-d4 timepoints did not have significant enrichment for any of the placental cell type clusters (Figure S7a,b). In contrast, the most mature iPS-derived TS cluster T7 was highly enriched for the syncytiotrophoblast (STB) and villous cytotrophoblast (VCT) cluster (p=log_10_^-32^ and log_10_^-17^) (Figure 5a). The expression profile of the iPS-derived TS T7 cluster was highly specific for STB and VCT with no significant enrichment for any other cell-type in the maternal fetal interface. The cluster BA4 (predominantly day 6 BMP4) was most enriched in the two fibroblast clusters (F1 and F2) (p=log_10_^-26^ and p=log_10_^-13^). Vento-Tormo described these placental fibroblasts as mesenchymal stromal cells of fetal origin that derive from the primitive endoderm expressing *GATA4*, *GATA6*, *PDGFRa*, and *SOX17* (Figure S7 and S8). The BI clusters, which contain mostly BMP4 + IWP2 treated cells, were not strongly enriched for specific placental cell type and instead had weaker enrichment for several clusters including syncytiotrophoblast (STB), and maternal decidua derived F1 endodermal cell type and dS1. In summary, iPS-derived TS cells are highly enriched for expression of genes specific to syncytiotrophoblast and villous cytotrophoblast of *ex vivo* human placenta.

**Figure 5:**
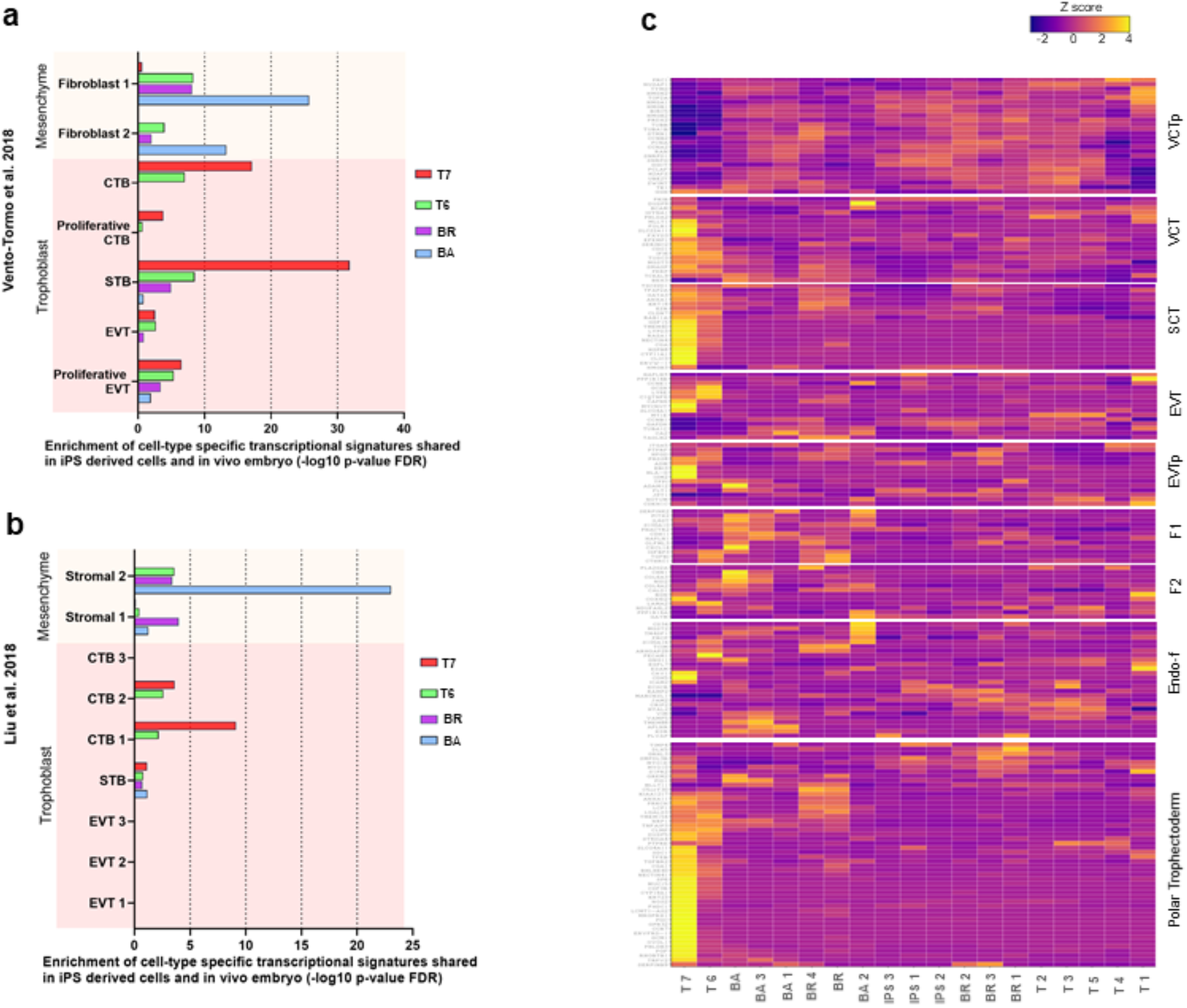
Trophoblast Differentiations Are Enriched For First Trimester Placental Cell Types. **(a,b).** Gene set enrichment analysis of most mature iPS-derived cell clusters as shown in figure 1c (BA4, BI5, T 6 and T 7n) to cell types identified in previous single cell sequencing of first trimester placenta identified in Vento-Tormo et al. 2018 [26] and Liu et al. 2018 [25]. **(c).** Composite heatmap of normalized gene expression of the top unique markers of placental cell types of fetal origin identified in Vento-Tormo et al. 2018 and polar trophectoderm markers (top 50) identified in Petropoulos et al. 2016 [44].

Next, we compared the expression profiles to an early placental dataset from the 8 week and 24 week placenta [25] (Figure 5b). Again, we found strong enrichment for the iPS-derived TS cells in the eight week fusion competent CTB (cytotrophoblast, log_10_^-9^). BA and BI clusters that contain mostly BMP4 and BMP4 + IWP2 conditions were enriched for two different mesoderm cells identified by Liu et al [25] (log ^-23^ and log_10_^-4^). To visualize placental cell-type specific expression patterns among the iPS-derived cell types, we plotted a heatmap of the top unique markers of the fetal cell types identified in Vento-Tormo and polar trophectoderm markers identified in Petropoulos [44] (Figure 5c). Consistent with the Gene Set Enrichment Analysis, we observed high expression for a majority of unique marker genes previously identified in polar trophectoderm villus cytotrophoblasts (VCT) and syncytiotrophoblasts (STB). We also compared the expression profiles of all iPS-derived cellular clusters to previously identified lineage and marker genes from pre-implantation datasets [45] (Figure S8a). Again, we found the highest expression of trophoblast genes *KRT18*, *TEAD3*, *GATA3* and *GATA2* in iPS-derived TS clusters, with moderate expression in BI clusters. Primitive endoderm genes *SOX17*, *GATA4*, *GATA6* were highest expressed in BA clusters. When compared to primary trophoblast cells (Figure S8b), we confirmed that the T7 cluster has the highest level of cytotrophoblast, trophectoderm and EVT enriched genes.

In summary, iPS cells specified via TS condition have a transcriptional profile highly similar to cytotrophoblasts taken directly from human placenta while the BMP4 differentiation conditions used here generates more heterogeneous placental cells types with evidence of amnion expression.

### Primed iPSC specify to trophoblast by activation of TFAP2A without activation of naive stem cell programs

It has been reported that primed stem cells are restricted in their potency and unable to differentiate to TSC while naive stem cells readily differentiate to TSC [41]. Therefore, we asked if the primed iPSC or TS condition cells adopt a naive stem cell program during differentiation. For 12 previously identified naive marker genes [46], naive transcripts were nearly absent. Five genes *HORMAD1*, *ALPPL2*, *KHD3L*, *TRIM60*, and *HYAL4* had no observed expression in any cell (Figure 6a, Table S7). Less than 1% of cells for any condition had detectable expression for six other marker genes *ALPP*, *OLAH*, *LYZ*, *MEG8*, *KDC1L*, and *FAM151A*. *ZNF729* was the only gene expressed in 0.51% of TSp1 cells to 14.69% of IWP2d2. In contrast, primed marker genes were consistently expressed in iPSC clusters and often throughout the differentiation state. For example, *DUSP6* was detected in 74.87% of the iPS cells, indicating a primed state. In sum, primed iPSC transitions to cytotrophoblasts without transitioning through a naive state.

**Figure 6:**
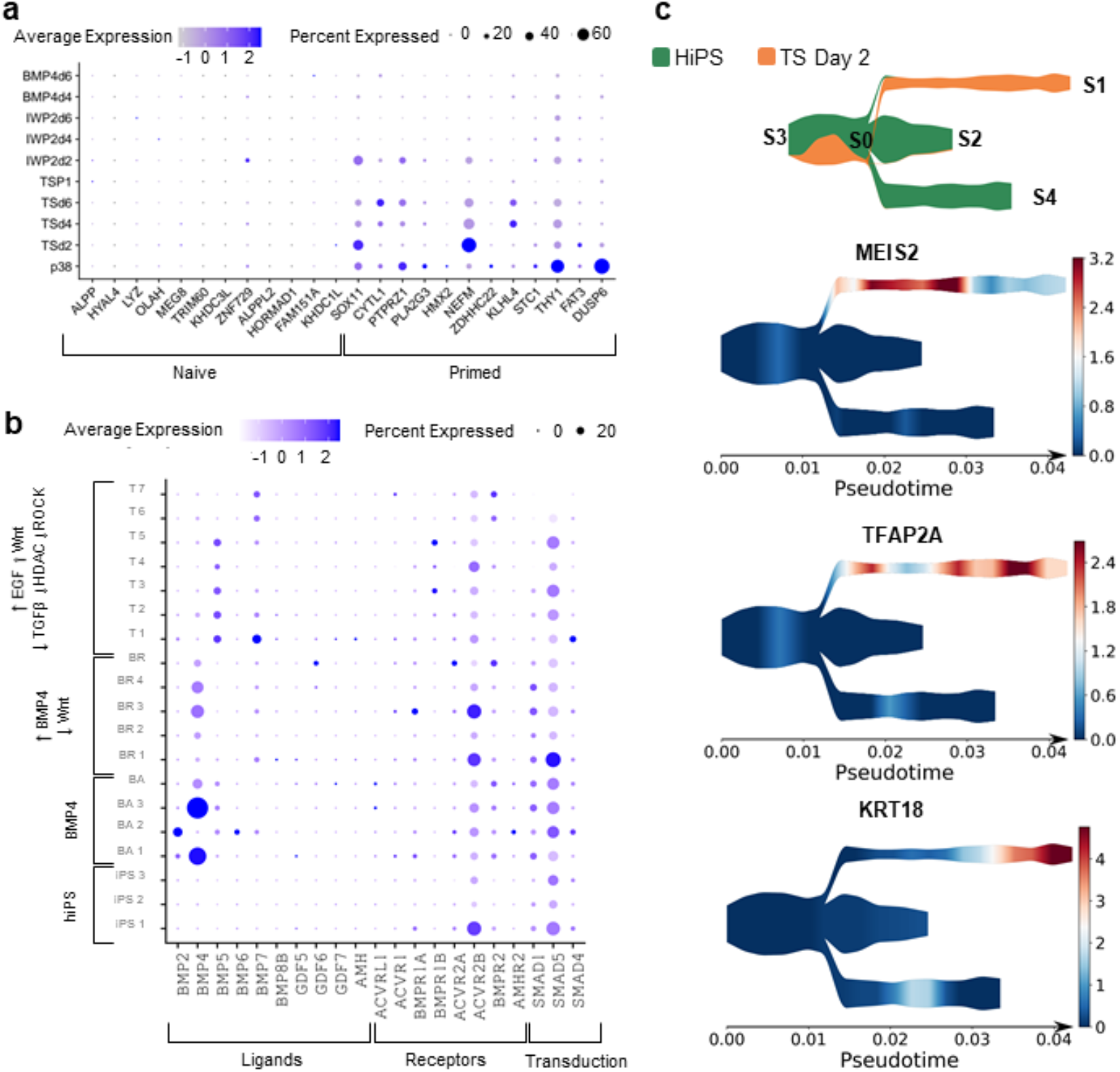
Primed iPSC Specification To Trophoblast Stem Cells Initiates With TFAP2A Rewiring Without Activation Of Naïve Stem Cell Programs. **(a).** Dotplot displaying expression of naïve and primed hPSC marker genes in the three differentiation treatments and days. See table S3 for source data containing log fold change and False discovery rate for each pairwise comparison by Wilcoxon Rank Sum Test with Holm-middle combined p value. **(b).** Dotplot displaying BMP signaling in each cellular cluster organized by the treatment identity for the majority of cells in the cluster. See table S3 for source data containing log fold change and False discovery rate for each pairwise comparison by Wilcoxon Rank Sum Test with Holm-middle combined p value. **(c).** STREAM pseudotime trajectory of hiPS differentiation for hiPS and TS D2 cells visualizing cells proportion along the smoothed pseudotime axis (orange represents TS, green represents hiPS). Nodes are labeled with numbers 0-4. Branches are defined as the cells between 2 nodes (Branch 1,4). Bottom: Top ranked (by STAT) genes with expression significantly correlated with the pseudotime transition of the branches. MEIS2, TFAP2A, and less differentiated TS d2 to more differentiated TSd2 (S0-S1) KRT18 (STAT 0.692). Ranks 2-4 for these branches is found in figure S6 c-d. Additional graphs showing the branch assignment with cells and a cell level representation of the pseudotime with the clusters figure S7 and table S7.

We asked how BMP signaling contributes to the differentiation by evaluating BMP family signaling in all samples. Low percentages of cells expressed BMP4 in TS condition, while BMP4 was significantly upregulated in several cell clusters of the BMP4 and BMP4 + IWP2 treated cells compared to iPSC and TS condition (Figure 6b, Table S7). BMP5 and BMP7 were expressed in a larger portion of TS condition cells compared to iPSC (BMP5: 0-0.67% iPSC vs 24.72% T1 cells and BMP7: 1.01-4.71% iPSC vs 29.06% T1 cells). The BMP effector proteins SMAD1,4, and 5 are expressed in the first days of all conditions, which can indicate that the endogenous BMP signaling may be important during the initial differentiation process.

To identify the most significant gene expression changes associated with the initial stages of TS condition specification, we applied a single-cell trajectory inference and pseudotime estimation (STREAM) [47] to the cells from the iPSC and TS condition day 2 samples. Briefly, single cells were ordered along probabilistic trajectories and a numeric value referred to as pseudotime was assigned to each cell to indicate how far it progresses along the dynamic differentiation. STREAM identifies an initial branch composed of both iPSC and TS condition cells at day 2 (S3-S0) which transition to three branches: TS condition day 2 (S0-S1) branch reflecting the exit from pluripotency, and iPS cell branches (S0-S2, S0-S4), reflecting the pluripotency continuum (Figure 6c and Table S8, S9, S10, S11, S12). We next identified the genes involved in the specification that are correlated with the transition along the branches (see methods). Pluripotency to TS (branch S0-S1) have upregulation of cytokeratins (*KRT18*, *KRT19*, *KRT8*) and calcium binding proteins (*S100A11*, *S100A10*) known to be expressed in trophectoderm [48, 49] (Figure S9d). Top positively correlated transition genes along the exit from pluripotency branch (S0-S1) include *LRRN3* which amplifies MAPK signaling through EGFR [50], *MEIS2* which codes for a TALE homeobox protein that promotes tumor progression through the combined action of two isoforms increasing YAP nuclear translocation and activating the WNT/b-catenin pathway [51], *BMP5*, *BMPR1B* and *TFAP2A* a transcription factor key to the suppression of pluripotency and expression of trophoblast associated genes [52] (Figure 6c, S9c). Also found as a top transition gene, *TMSB4X* is implicated in stemness of progenitor trophoblast cells of first trimester human placentas by increasing NOTCH1 activity [53, 54, 55]. In summary, trophectoderm transcriptional programming occurs early in the specification and utilizes previously identified trophectoderm associated regulatory factors including *TMSB4X*, *YAP*, *BMP5*, and *TFAP2A*.

### Human endogenous retrovirus derived genes participate in a regulatory subnetwork within primed-derived trophoblast stem cells

Endogenous retroviruses play a fundamental role in the evolutionary diversification of the mammalian placenta with multiple instances of human endogenous retroviruses (hERVs) co-option [56, 57, 58, 59]. We reasoned that hERV expression may be altered because primed PSC, naive PSC and TSC differ in the regulation of human endogenous retroviruses [60], and therefore, we investigated how hERVs are expressed during specification. While there are potentially 1500 ERV-derived genes capable of encoding proteins [57], we restrict our analysis to the 20 ERV-derived genes that are currently annotated as human genes including *ERV3-1, Syppressyn (ERVH48-1)*, *syncytin-1 (ERVW-1)* and *syncytin-2 (ERVFRD-1),* which function in antiviral responses and syncytial fusion. We found specific upregulation of five ERV-derived genes (*ERVH48-1*, *ERV3-1*, *ERVMER34-1*, *ERVW-1* and *ERVFRD-1*) in the most differentiated cells in the TS condition, cluster T7 (Figure S10a).

To understand the place of ERV-derived genes in the regulatory TS regulatory network, we analyzed the participation of ERV-derived genes in a gene regulatory network. We built a transcription factor and target gene network model using the Passing Attributes between Networks for Data Assimilation (PANDA) algorithm to all clusters among the different conditions used here. PANDA integrates information from TF-sequence-motif data, gene expression and protein-protein interaction (PPI) in a message-passing approach [61]. We found that ERV interactions (Transcription Factors + ERV-derived genes) are prominent in the most differentiated TS cluster TS 7 and in no other cell clusters. We found regulatory interactions for five ERV-derived genes (*ERVH48-1*, *ERV3-1*, *ERVMER34-1*, *ERVW-1* and *ERVFRD-1*) are highly unique to the TS 7 cluster network, while only 0 or 1 ERV-derived gene was found in the other cell clusters (Figure S10b, S11a). Interestingly, genes encompassed in ERV regulatory interactions of primed-derived TSC were enriched in biological pathways related to hormone metabolism, cell differentiation and the immune system (adj. p value < 0.01) (Figure S10c, Table S13). In addition, ERV regulatory interactions are also enriched for categories such as placenta development, trophectodermal cell differentiation and syncytium formation (Table S13). To understand the significance of the ERV-subnetwork interactions, we next asked what is the strength of ERV edges in the network context of the TS cluster. Weights of ERV regulatory interactions were highly ranked among all network connections (Wilcoxon Rank Sum p-value=1.6778e^-31^, permuted p-value=0.001), indicating a significant contribution to the gene regulatory network of the TSC state. In sum, we find that ERV-derived genes participate in a regulatory subnetwork within primed-derived TSC.

## Discussion

Here we map the specification of primed iPSC to trophoblast by exogenous induction of EGF, WNT and Valproic acid and inhibition of TGF-β and HDAC and demonstrate the involvement of GATA3, BMP5/7, TFAP2A without transitioning through a naive state. Primed-derived-TSC are highly transcriptionally similar to cytotrophoblasts found in human placenta. Primed-derived-TSCs proliferate rapidly as a stem cell and differentiate into multinucleated extravillous and syncytiotrophoblast. After eight days of specification, primed-derived-TSC displays a uniquely active transcriptional network of human endogenous retroviruses similar to *in vivo* trophoblast.

The ability of primed human stem cells to differentiate into true trophectoderm lineages has been contested. Here we confirm that primed PSC retains a broad potency to generate extra embryonic lineages which are transcriptionally highly similar to placenta. While recent studies conclusively prove that primed PSC can be specified to TSC, studies report absent [62] or inefficient conversion to TSC with the direct application of the TSC-media, as described here. Optimized culturing conditions and differences in the primed PSC state at the initiation of TSC-media exposure likely explain the more robust differentiation described here. We induce differentiation after self-organized multicellular colonies are established by clump-passaging primed cells from StemFlex conditions, plating cells in StemFlex media and inducing differentiation 24hrs after passaging all in 20% O2. Instead of StemFlex, others have initiated differentiation with primed PSC cultured on MEFs (Viukov et al., 2022), TSER or E8, a xeno-free media, and used either 20% or 5% O2. StemFlex is proprietary media containing bovine serum albumin and not all components are known. While PSCs grown in StemFlex and E8 have similar transcriptional profiles, PSC in StemFlex demonstrate less karyotypic abnormalities, better expression of Nanog, more efficient single cell cloning functional differences such as enhanced single-cell cloning and robustness to automation (Daniszewski et al., 2018). While the conditions optimized here improve the specification of TS from primed iPS, removing WNT for the initial days would likely improve the efficiency even further, as recently published for primed PSC grown on MEFS [10].

We also confirm that the exogenous BMP4 protocols can generate cells with key features of trophoblast, but the majority of cells specified by BMP4 are less similar to *in vivo* cytotrophoblasts. Importantly, the BMP4 methods used here differ from the more established BAP method, which are optimized to produce more efficient trophectoderm differentiation. In those optimized conditions, transient BMP4 application induces a naive state. For the direct TSCM method here, we find that endogenous BMP5 and BMP7 are induced, without inducing BMP4 or naive state. Similar to induction with BMP4, direct TSCM method induces transient initial CDX2 expression, and CDX2 is also transiently upregulated in an enhanced primed-to TSC conversion without WNT stimulation and inhibiting TGF-B.

Our data suggest that while PSC can differentiate via multiple paths, the paths share many established trophectoderm specification factors and the final TSCs are similar. Single cell sequencing allows deconvolution of the cellular specification by complete transcriptional profiles. The specification generates two separate cell clusters both with trophectoderm expression, and the primed-derived TSCs preferentially expand to purity with passaging (Figure 1p). Further supporting similarity to primary TSC derived from human villous cytotrophoblast and from the blastocyst outgrowth, the primed-derived-TSC also begin as mixed populations of cells which are passaged to generate pure TSC lines [3]. Our trajectory analysis indicates that *CDX2* and *TFAP2C/A* are important in the initial exit from pluripotency in primed cells and facilitates trophectoderm lineage specification, similar to human trophectoderm specification *in vivo.* TFAP2A is a transcription factor involved in both trophoblast specification and a trophoblast-like amnion speciation. In humans, the first wave of amnion specification may follow a trophectoderm-like pathway [11]. We use multiple statistical tests examining single genes, gene sets and complete transcriptional profile similarities to demonstrate that the primed-derived-TSC are most transcriptionally similar to trophectoderm and not amnion.

We find that five hERV-derived ORFs annotated as human genes are expressed in primed-derived TSC, similar to in vivo [57]. hERV are abundant sequences within the human genome that are derived from ancient retroviral infections in human ancestral germline. hERV-derived Long Terminal Repeats function as human-specific enhancers during placentation and inflammation. hERV-derived envelope proteins create coding ORFs. A specific hERV-envelope protein named syncytian-1 (hERVW-1) is best characterized for cell-cell fusion during embryo attachment and syncytialization where it is upregulated [58]. Like many of the other 668 hERV-envelope proteins expressed in the early embryo and placenta, Syncytian-1 is widely expressed in ICM, trophectoderm and CTB [57]. Other hERV-derived ORFs inhibit cell fusion (syncytin-2), provide immunosuppressive activities (syncytin-2, ERV3-1 and suppressyn) [59], and restrict retroviral infection in morula, ICM, TE and ESC (supressyn) [57].

The differentiation of primed iPSC to trophectoderm allows for modeling of placental diseases from patient-specific iPS cells without first trimester primary tissue and without transitioning through a naïve state. While the final TS product of these different models appears similar, the naive vs primed states differ in molecular features which could later impact hTSC models. In naive state, global DNA demethylation and genomic instability occur in both t2i/L/Go or 5i/L/A conditions [64, 60], and imprinted loci may be aberrantly demethylated [40]. Mammalian imprinting is most pronounced in the placenta [66] and cellular conditions that maintain imprinting may be important in a cellular model of human placenta. X chromosome inactivation (XCI) is reset in naive conditions compared to inactivated or eroded in primed conditions, and it remains to be seen if there are differences in XCI.

Placental adaptive responses to insults can alter fetal development and program the fetus for later disease including diabetes and neurodevelopmental disorders [67, 68]. As both the protective barrier and a component of the innate immune system, cytotrophoblasts and their progeny play important roles in the outcome of insults [69]. Recent evidence that Covid19 leads to a hyperimmune state has implications for placental biology and fetal development, potentially analogous to the effects of other hyperimmune states during pregnancy [2]. Genetic variation among individuals influences how a pathogen affects the fetus. iPSC models of this genetic variation could be harnessed for therapeutic intervention. We anticipate that primed-derived TS cells will be a powerful model to study the placental contribution to disease and the genetic regulation of human placental evolution.

## Material and Methods

### iPSC culture

Human ES and iPSC were maintained on Collagen IV (1mg/12 ml DMEM/F12) coated plates in StemFlex media. Media was changed every 48hrs in accordance with manufacturer recommendations. Cells were passaged in small clusters using Versene solution. See supplemental methods for details.

### Differentiations and Cell Maintenance

24 hours post passaging with Versene, cells were washed with DPBS and media was changed to differentiation media. TS medium: [DMEM/F12 with Glutamax supplemented with 0.1 mM 2-mercaptoethanol, 0.2% FBS, 0.3% BSA, 1% ITS-X supplement, 1.5 μg/ml L-ascorbic acid, 50 ng/ml EGF, 2 μM CHIR99021, 0.5 μM A83-01, 1 μM SB431542, 0.8 mM VPA and 5 μM Y27632]. BMP4: 10 ng/ml BMP4 with/without 2uM IWP2 was added to basal differentiation medium (DMEM/F12 with Glutamax supplemented with ITS and L-ascorbic acid 2-phosphate magnesium). Cells were collected on day 2, 4, and 6 for RNA sequencing. For the TS condition, after 6 days of differentiation cells were passaged using TrypLE express at a split ratio of 1:3 and plated on new Collagen IV (5 ug/mL) coated plates. After the first passage, media was changed every 48 hours and split at a ratio of 1:3-1:6 every 3-4 days. See supplemental methods for more details.

### EVT and STB differentiation and characterization

Differentiation of TSCs were performed as previously described [3], with minor modifications. For EVT differentiation, 24-well plates ibidi were coated with 1 μg/mL Collagen IV overnight. 2 × 10^5^ TSC were seeded per well in 500µL EVT basal medium [DMEM/F12 supplemented with 0.1 mM β-mercaptoethanol, 0.5% penicillin-streptomycin, 0.3% BSA, 1% ITS-X, 7.5 μM A83-01, 2.5 μM Y27632] supplemented with 4% KSR and 100 ng/mL NRG1 alpha. Matrigel was added to a 2% final concentration shortly after resuspending TSC in the medium. On day 3, the media were replaced with 500µL EVT basal medium supplemented with 4% KSR, and Matrigel was added to a 0.5% final concentration. At day 6, the media were replaced with 500µL EVT basal medium, and Matrigel was added to a 0.5% final concentration. At day 9, the cells were ready for analysis

For STB differentiation, 24-well plates ibidi were coated with 2.5 μg/mL Collagen IV overnight. 2 × 10^5^ TSCs were seeded per well in 500µl STB medium [DMEM/F12 supplemented with 0.1 mM β-mercaptoethanol, 0.5% penicillin-streptomycin, 0.3% BSA, 1% ITS-X, 2.5 μM Y-27632, 2 μM Forskolin and 4% KSR]. The media was changed at day 3, and at day 4 the cells were ready for analysis.

### Data Availability

The Single cell RNA sequencing datasets generated and/or analyzed during the current study have been deposited in ArrayExpress with the accession code E-MTAB-9526. Analysis pipelines are publicly deposited on Github:https://github.com/paquolalab/placenta_ips

## Supporting information

Supplemental Figures, Methods, Tables, Key Reagents and Resources

## Acknowledgements

Supported by funding from the Lieber Institute for Brain Development and a NARSAD Young Investigator Grant from the Brain & Behavior Research Foundation to J.A.E, the National Institute of Health (NIH) T32 Fellowship KJB (T32MH015330). The authors are grateful for the financial support of the Coordenação de Aperfeiçoamento de Pessoal de Nível Superior (CAPES) - Financing code 001 (Andre Barbosa, Veronica Euclydes and Arthur Feltrin). This work was supported in part by FAPESP - Sao Paulo Research Foundation (http://www.fapesp.br/en/): Grant Number 2018/18560-6. We thank Bill Ulrich for the single cell RNAseq browser. We thank Tricia Nilles and Worod Allak from the Becton Dickinson Immune Function Laboratory at the Johns Hopkins Bloomberg School of Public Health, for flow cytometry technical assistance. The facility was supported in part by CFAR: P30AI094189-04 (Chaisson).

## Author Contributions

JE and ET. conceptualized the study; JE, ET, ARB, BHSA and VE. wrote the original draft; JE, ET, ARB, BHSA, VE, HJC, YKL, AF, JL, PDC, TS, and KJB performed experiments and data analysis; HB, JEK, TMH, BGS, DRW, GU, RM, JHS, ACMP, JE were responsible for writing, reviewing, and editing of the manuscript; JE coordinated the study.

## Classification

Biological Science and Developmental Biology.

