## Supplemental Figures, Methods, Tables, Key Reagents and Resources for "A single cell trajectory of human archetypal pluripotent stem cell differentiation to trophoblast stem cells reveals induction of endogenous BMP5/7 and GATA3 without transitioning through a naive state"

[#]Equal contributor

**This PDF file includes:**

Supplementary text
Figures S1 to S11
Tables S1 to S8
Key Reagents and Resources
Supplemental Methods

**Supplemental Figures and Legends**

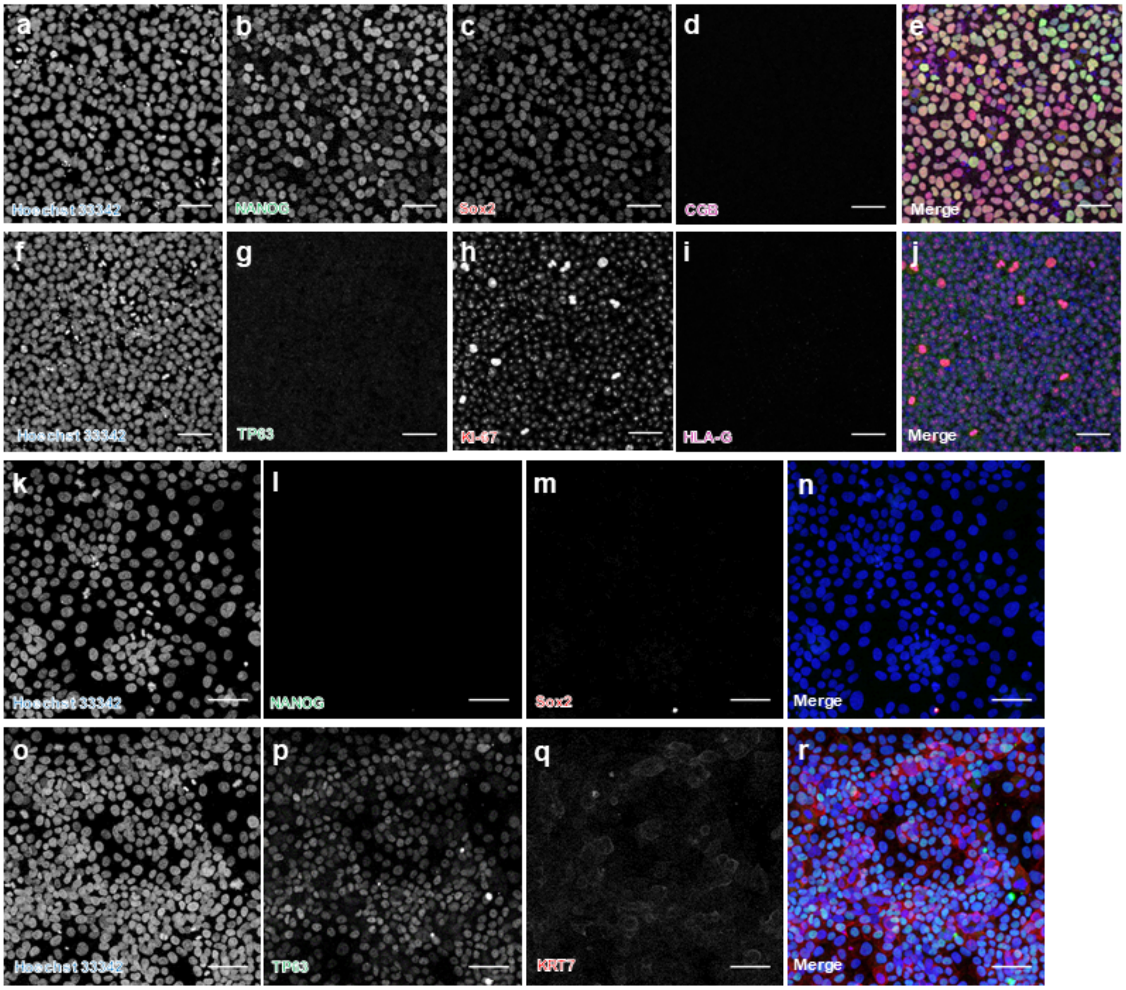

**A.** Boxplot of Mitochondrial Mapping rate per droplet in the final data set.

**B.** Boxplot of number of detected genes per droplet in the final data set.

**C-D.** Summary tables of cell number and percent composition by experimental sample (C) and cluster identity (D).

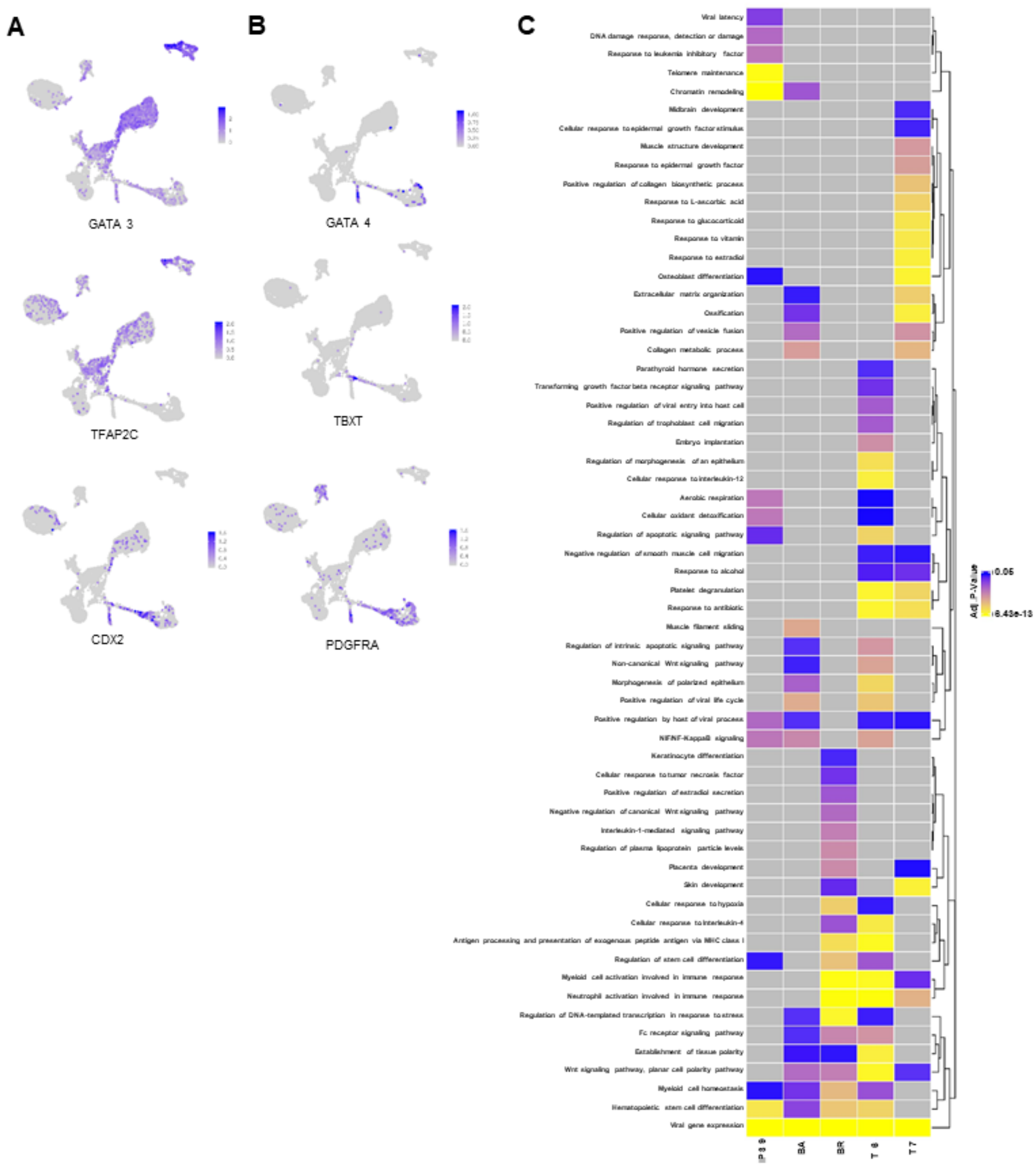

**Fig. S2: BMP4 model comparison.**

**A-B.** Highlighted expression of trophoblast (A) and mesoderm (B) associated transcription factors at single cell resolution.

**C.** Heatmap of selected GO terms (p-value <0.05) generated from the 300 most highly expressed genes for each of the indicated mature cell types.

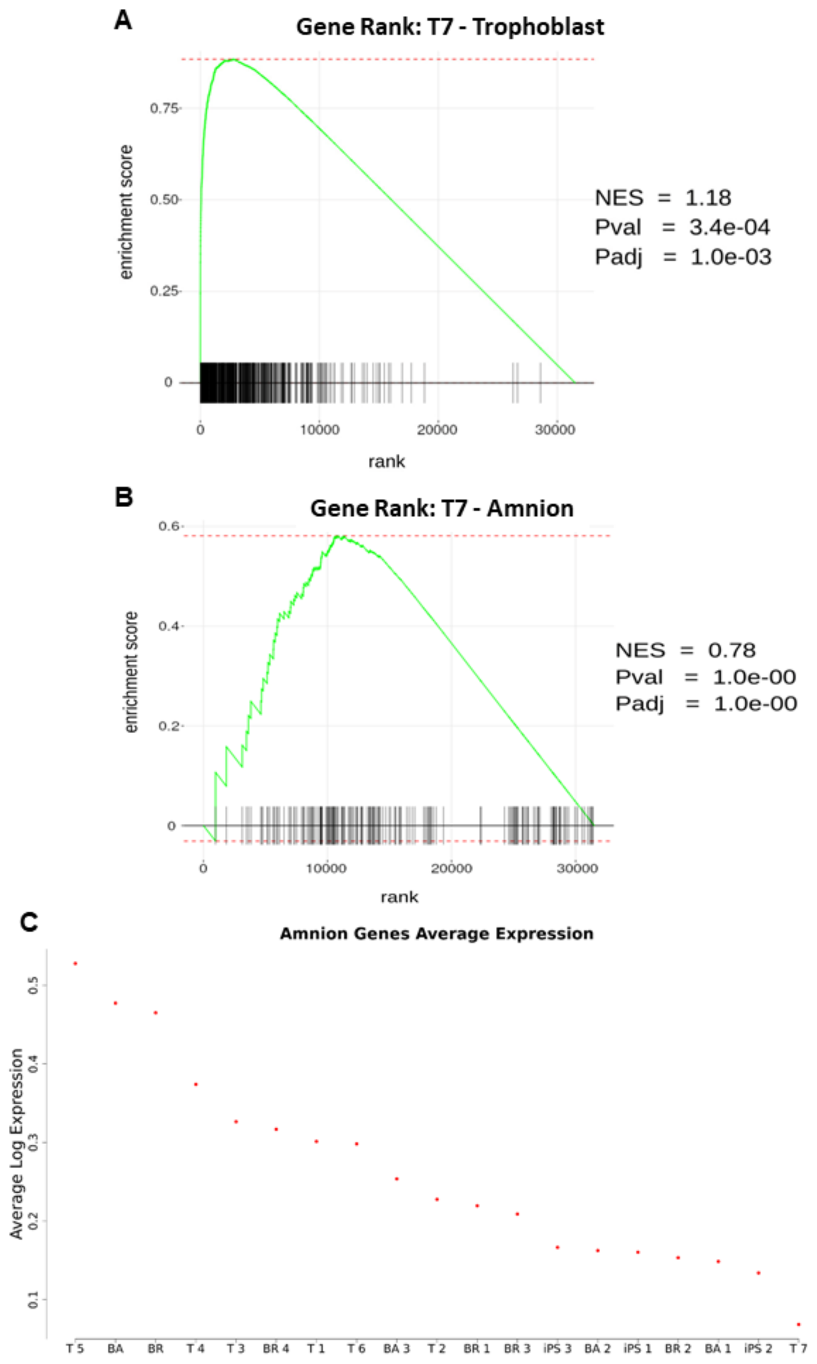

**Fig. S3: Gene Set Enrichment Analysis for Amnion and Trophoblast genes.**

**A-B**. Gene rank for trophoblast and amnion in T7 sample.

**C.** Mean expression of amnion genes among all clusters.

A

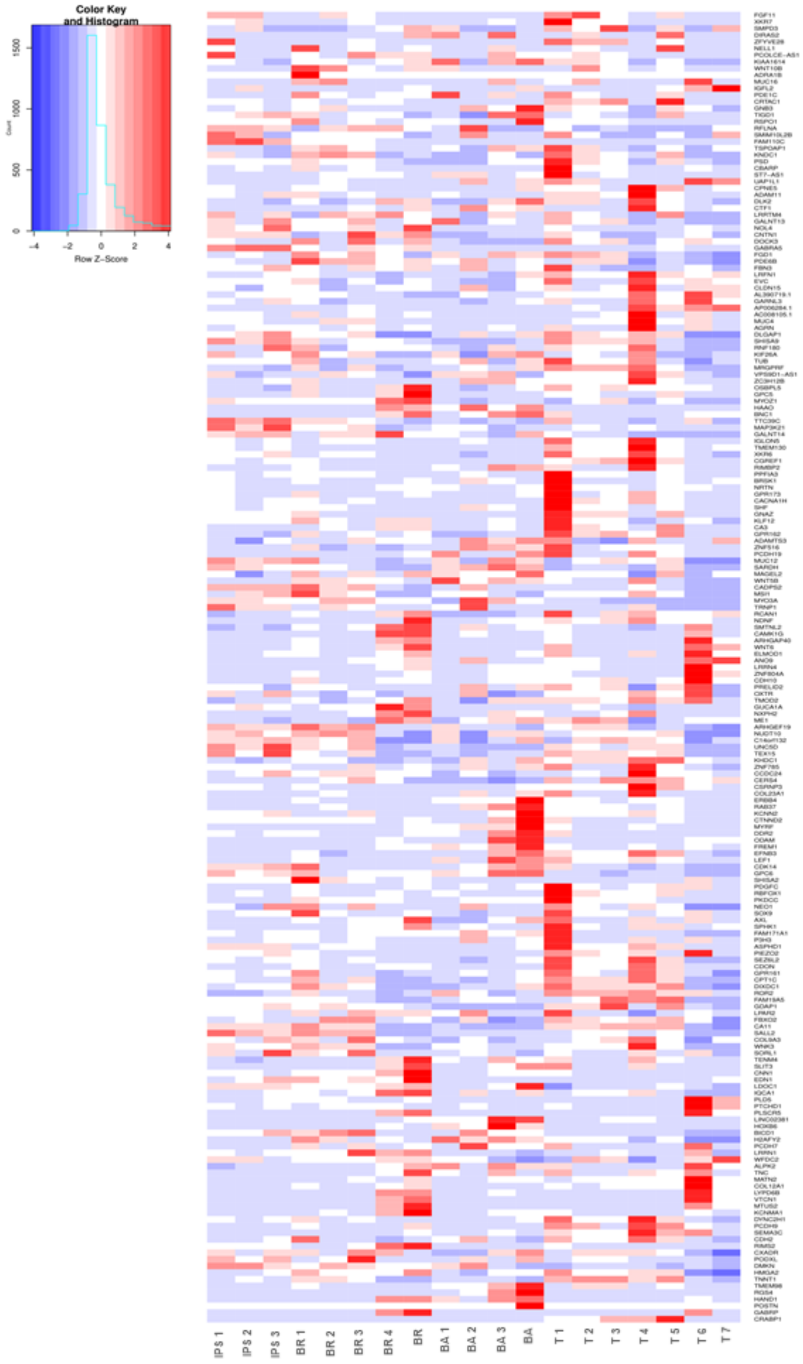

**Fig. S4. Heat map of human amnion genes**.

**A.** Heat map of the top genes predominantly expressed in human amnion [1] in each cluster.

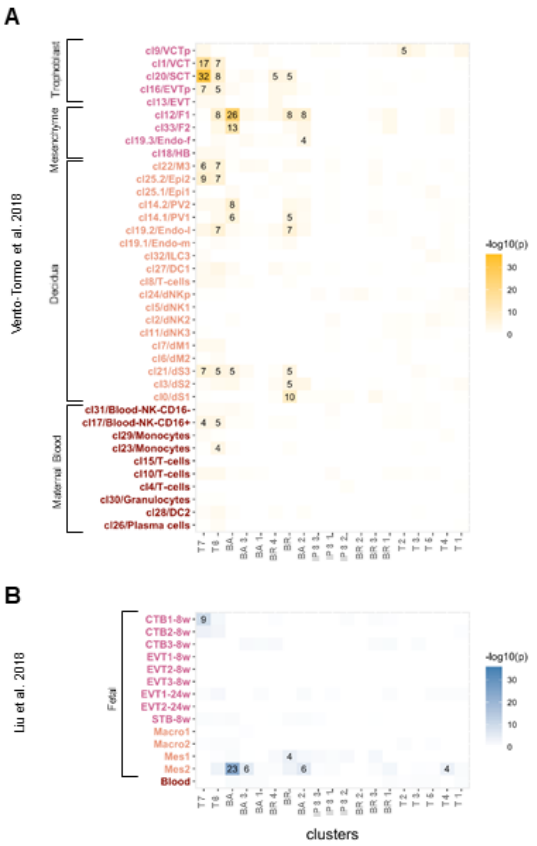

**Fig. S5: Trophoblast differentiations are enriched for first trimester placental cell types.**

**A,B.** Gene set enrichment analysis of HiPS cell differentiations highlighting enrichment of cell clusters to cell types identified in first trimester placentas. Color and number in each square indicates the –log10(p) Bonferroni Corrected hypergeometric test for enrichment of cell-specific genes in IPS-derived clusters and previous placental publications.

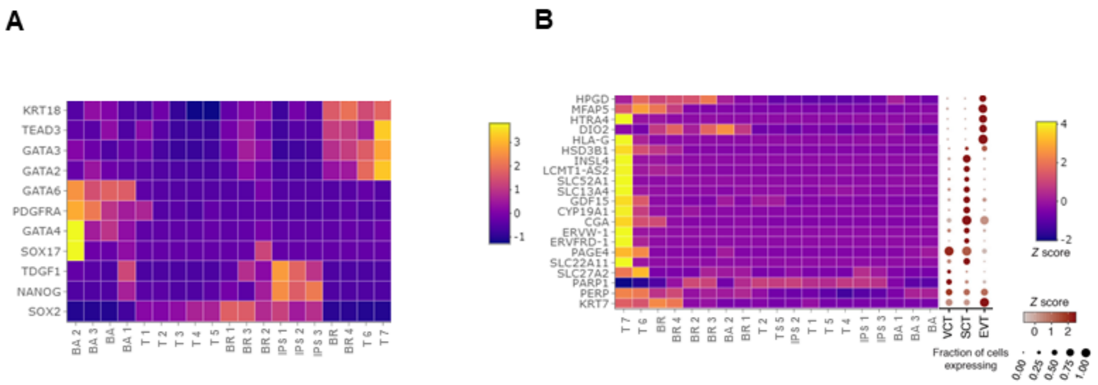

**Fig. S6: Lineage specification and cellular cluster identity based on marker gene profiles.**

**A.** Heatmap of detected high confidence markers for epiblast, primitive endoderm, and trophectoderm from Stirparo et al. 2018 [2].

**B.** Heatmap of trophoblast specific markers from Suryawanshi et al. 2018 including modified dotplot from their publication for primary Villous Cytotrophoblasts (VCT), Extravillous Trophoblasts (EVT), and Syncytiotrophoblast (SCT).

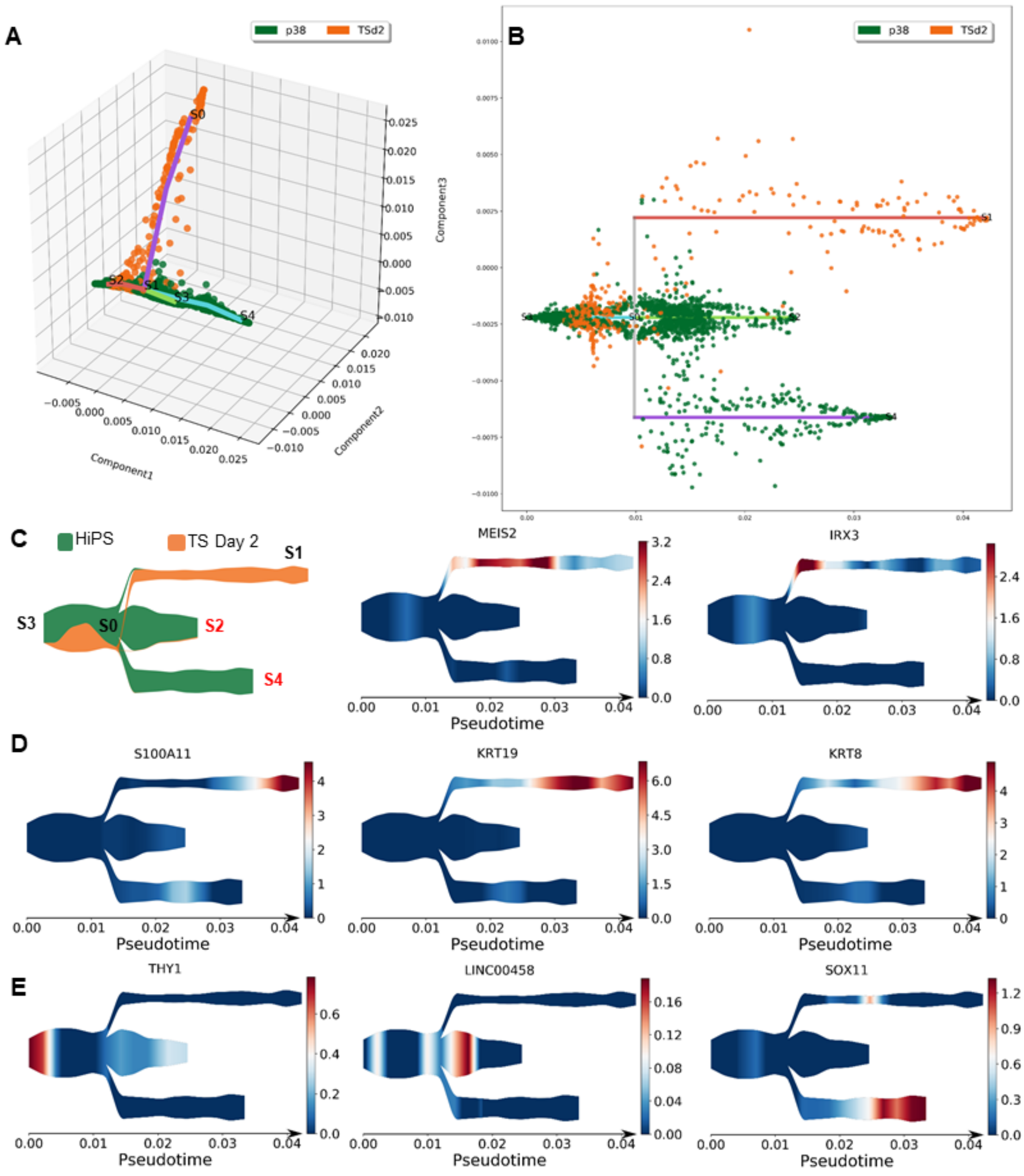

**Fig. S7: Developmental trajectory of HiPS and after two days of differentiation in TS media.**

**A-B.** Visualization of branch assignment using the first 3 SE components (A) and resulting subway map representation of the reconstructed pseudotime trajectory using HiPS and TS D2 cells samples (B).

**C-D.** Averaged expression of significant transition genes along the pseudotime trajectory ordered left to right by rank (2-4) for branch S3-S0 (C) and S0-S1 (D).

**E.** Averaged expression of selected primed pluripotency markers expressed along psuedotime.

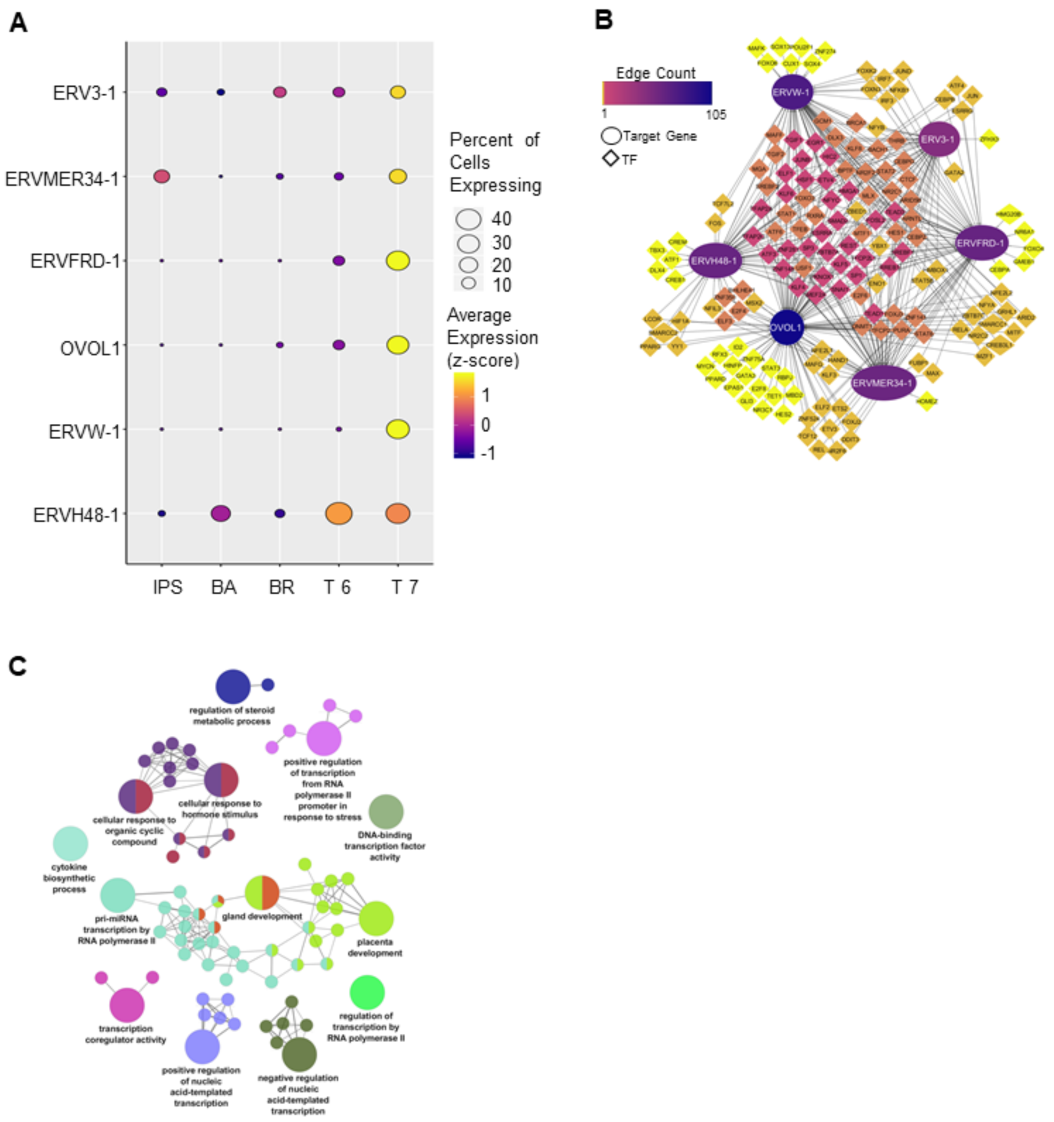

**Fig. S8: Endogenous Retrovirus-derived Genes Participate In A Gene Regulatory Subnetwork Uniquely In TS Cells.**

**A**. Dotplot of HERV expression in most mature cell clusters from each treatment and the HiPS cluster 1. Size of the ball indicates the percent of cells expressing the gene and color represents the z-normalized expression value.

**B**. Cytoscape representation of the top transcription factor partners of ERVs detected in TS and BA. 164 TFs and 6 Targets. See Supplemental table S8.

**C**. Visual Representation of GO terms from most significant partners of ERVs shown in panel B (p<.0001). Specifics of programs and settings to obtain these results. Cytoscape 3.7.2, cluego v2.5.4, cluepedia version v1.5.4, go term fusion, levels 3-8, kappa .35, graph of highest significance within each group, input of 164 TFs from panel B supplemental table with analysis output and details of GO analysis includes all go terms.

**A**

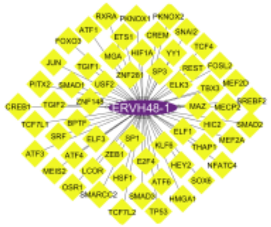

**Fig. S9: PANDA network analysis reveals unique regulation of endogenous Retroviruses in TS cells.**

**A.** ERV48-1 connected network identified by PANDA in BA cells.

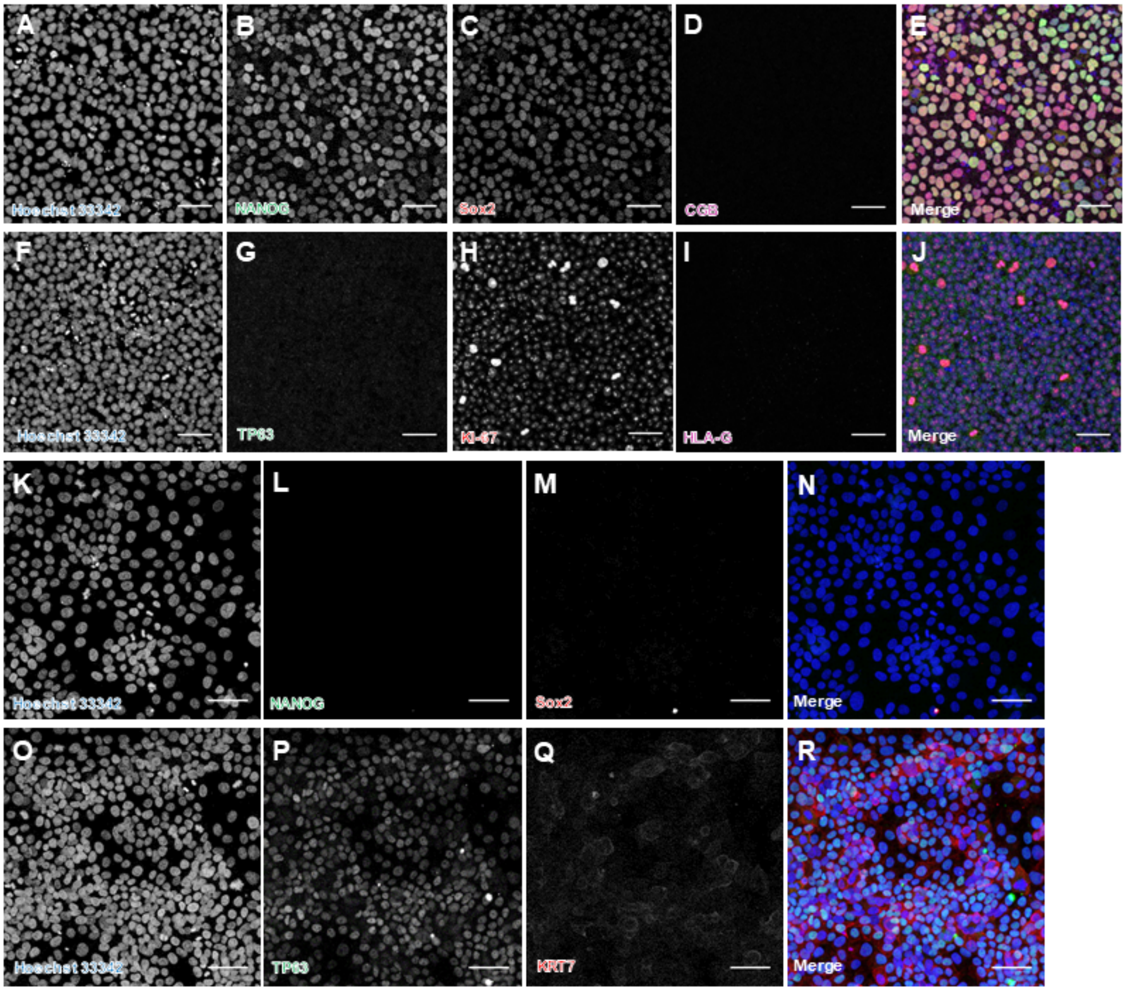

**Fig. S10: Differentiated TS cells lose expression of pluripotency markers while gaining expression of trophoblast specific markers.**

**A-E**. Immunofluorescent images of HiPS line LIBD7C6 stained for Hoechst 33342 (A), NANOG (B), and Sox2 (C), CGB (D), and in composite (E).

**F-J**. Immunofluorescent images of HiPS line LIBD7C6 stained for Hoechst 33342 (F), TP63(G), and KI-67 (H), HLA-G (I), and in composite (J).

**K-N**. Immunofluorescent images of TS line 2014.06 stained for Hoechst 33342 (K), NANOG(L), and Sox2 (M), and in composite (N).

**O-R**. Immunofluorescent images of TS line 2014.06 stained for Hoechst 33342 (O), TP63 (P), KRT7 (Q), and in composite (R).

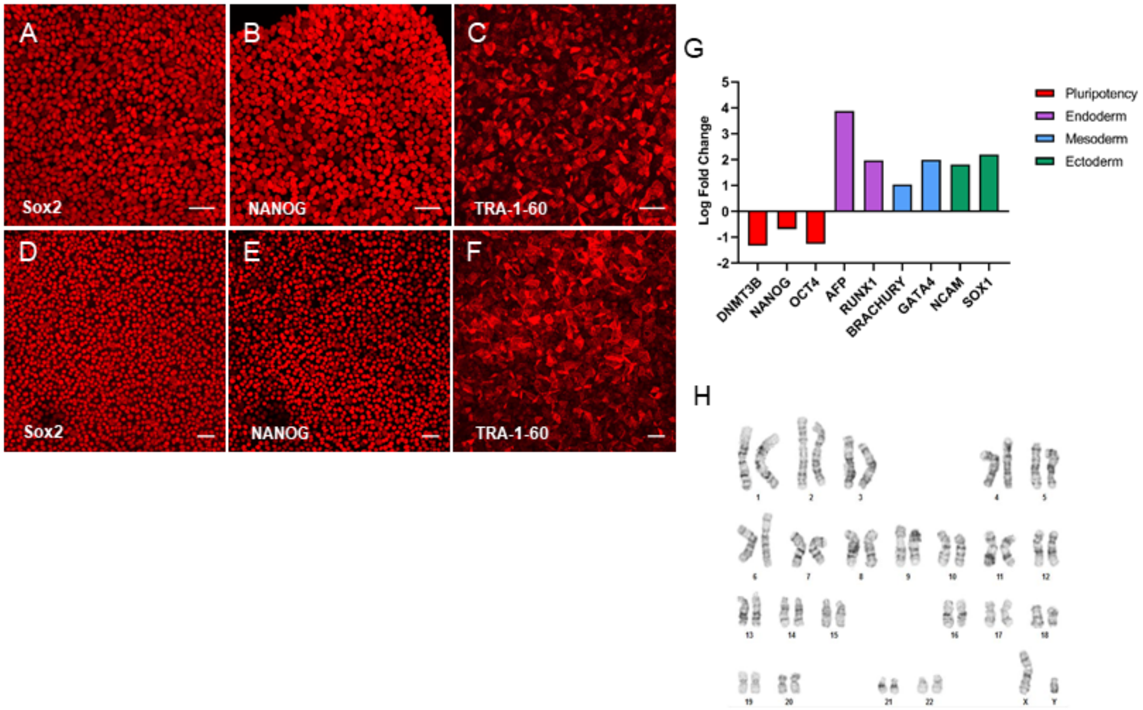

**Fig. S11: iPS cell line characterization.**

**A-C**. Immunofluorescent images of HiPS line LIBD1C8 stained for Sox2 (A), NANOG (B), and TRA-1-60 (C).

**D-F**. Immunofluorescent images of HiPS line 2014.06 stained for Sox2 (D), NANOG (E), and TRA-1-60 (F).

**G**. Relative expression levels of pluripotency and germ layer specific markers in spontaneously differentiated cells of line 2014.06 normalized to their respective expression levels in undifferentiated HiPS of the same cell line determined by qPCR.

**H**. Karyotype analysis of 2014.06

All scale bars 50 um.

**Table S1 (separate file).** Number of cells from each condition/day that are assigned to the cluster

**Table S2 (separate file).** Differentially expressed genes in each cluster where a single cluster is compared to all other clusters

**Table S3 (separate file).** Differentially expressed genes analyzed by pairwise comparisons of one cluster compared to another cluster

**Table S4 (separate file).** Average log normalized expression of amnion genes [1] for each clusters.

**Table S5 (separate file).** Expression analysis of naive, primed and BMP signaling during differentiation

**Table S6 (separate file).** Stream pseudotime analysis of transition and differentially expressed leaf genes for iPS to TSd2

**Table S7 (separate file).** Stream pseudotime analysis of transition and differentially expressed leaf genes for all cells

**Table S8 (separate file).** ERV gene regulatory network and associated GO enrichment.

**Key Reagents and Resources**

| Reagent or Resource Type | Name | Supplier | Product Number | Additional information |
|---|---|---|---|---|
| Antibody | anti-hCGβ (mouse monoclonal) | Abcam | ab9582 | 1:100 |
| Antibody | anti-HLAG (mouse monoclonal) | Abcam | ab52455 | 1:100 |
| Antibody | anti-KI67 (rabbit polyclonal) | Abcam | ab15580 | 1:250 |
| Antibody | anti-KRT7 (mouse monoclonal) | ThermoFisher Scientific | MA5-11986 | 1:1000 |
| Antibody | anti-TJP1 (mouse monoclonal) | ThermoFisher Scientific | 33-9100 | 1:100 |
| Antibody | anti-TP63 (goat polyclonal) | Novus Biologicals | AF1916 | 1:100 |
| Antibody | anti-VIM (rabbit monoclonal) | Abcam | ab92547 | 1:500 |
| Antibody | anti-SDCA1 (rabbit monoclonal) | Abcam | Ab128936 | 1:500 |
| Antibody | anti-CD49f (Integrin alpha 6 – rat monoclonal - FITC) | eBioscience | 11-0495-82 | 0.25μg |
| Antibody | Isotype control anti-IgG2a kappa (rat – FITC) | eBioscience | 11-4321-42 | 0.25μg |
| Antibody | Donkey anti-mouse IgG AF-488 | Jackson Immunoresearch Laboratories | 715-545-151 | 1:500 |
| Antibody | Donkey anti-mouse IgG Cy3 | Jackson Immunoresearch Laboratories | 715-165-151 | 1:500 |
| Antibody | Donkey anti-rabbit IgG AF-488 | Jackson Immunoresearch Laboratories | 711-545-152 | 1:500 |
| Antibody | Donkey anti-rabbit IgG Cy3 | Jackson Immunoresearch Laboratories | 711-005-152 | 1:500 |
| Antibody | Donkey anti-rabbit IgG AF-647 | Jackson Immunoresearch Laboratories | 711-605-152 | 1:500 |
| Antibody | Donkey anti-goat IgG Cy3 | Jackson Immunoresearch Laboratories | 705-165-147 | 1:500 |
| Chemical, Peptide, or Recombinant Protein | 2-mercaptoethanol | ThermoFisher Scientific | 21985023 | |
| Chemical, Peptide, or Recombinant Protein | Bmp4 | Peprotech | 120-05-5ug | |

| | | | |
|---|---|---|---|
| Chemical, Peptide, or Recombinant Protein | Bovine Serum Albumin | Cell Signaling Technology | 9998S |
| Chemical, Peptide, or Recombinant Protein | Cell Banker 1 Cryopreservation Media | Amsbio | 11888 |
| Chemical, Peptide, or Recombinant Protein | Collagen IV | Corning | 354233 |
| Chemical, Peptide, or Recombinant Protein | Cultrex Reduced Growth Factor Basement Membrane Extract, PathClear | R&D Systems | 3433-005-01 |
| Chemical, Peptide, or Recombinant Protein | DMEM/F12, Glutamax supplement | ThermoFisher Scientific | 10565018 |
| Chemical, Peptide, or Recombinant Protein | DPBS, no calcium, no magnesium | ThermoFisher Scientific | 14190250 |
| Chemical, Peptide, or Recombinant Protein | Epidermal Growth Factor | Peprotech | AF-100-15 |
| Chemical, Peptide, or Recombinant Protein | Fetal Bovine Serum | ThermoFisher Scientific | 16140071 |
| Chemical, Peptide, or Recombinant Protein | Gelatin | Sigma-Aldrich | G1393-100mL |
| Chemical, Peptide, or Recombinant Protein | Hoechst 33342 | ThermoFisher Scientific | H3570 |
| Chemical, Peptide, or Recombinant Protein | Insulin-transferrin-selenium | Corning | 25-800-CR |
| Chemical, Peptide, or Recombinant Protein | ITS-X | ThermoFisher Scientific | 51500056 |
| Chemical, Peptide, or Recombinant Protein | IWP2 | BioGems | 6866167-1mg |
| Chemical, Peptide, or Recombinant Protein | L-Ascorbic Acid | BioGems | 2520691 |
| Chemical, Peptide, or Recombinant Protein | L-ascorbic acid 2-phosphate magnesium | Sigma-Aldrich | A8960-5G |
| Chemical, Peptide, or Recombinant Protein | SB431542 | BioGems | 3014193 |
| Chemical, Peptide, or Recombinant Protein | StemFlex Media | ThermoFisher Scientific | A3349401 |
| Chemical, Peptide, or Recombinant Protein | TrypLE express | ThermoFisher Scientific | 12604013 |
| Chemical, Peptide, or Recombinant Protein | Valproic Acid Sodium salt | BioGems | 1066656 |
| Chemical, Peptide, or Recombinant Protein | Y-27632 | BioGems | 1293823 |
| Chemical, Peptide, or Recombinant Protein | Forskolin | BioGems | 6652995 |

| | | | | |
|---|---|---|---|---|
| Chemical, Peptide, or Recombinant Protein | Recombinant human NRG1 alpha | BioLegend | 765704 | |
| Cell Line | 2014.06 Human Induced Pluripotent Stem Cells | Lieber Institute | N/A | |
| Cell Line | CF-1 Mouse Embryonic Fibroblasts | ThermoFisher Scientific | A34181 | |
| Cell Line | LIBD1C8 Human Induced Pluripotent Stem Cells | Sawada et al., 2020 | N/A | Generated in lab |
| Cell Line | LIBD7C6 Human Induced Pluripotent Stem Cells | Sawada et al., 2020 | N/A | Generated in lab |
| Cell Line | WA-01 (H1) Human Embryonic Stem Cells | Wicell | WB0197 | |

**Supplemental Methods**

***PSC culture:*** 6-well plates were coated with reduced growth factor Cultrex (1mg/12 ml DMEM/F12) at 37°C for at least one hour. Human ES and iPS were maintained on coated plates in Stemflex media. Media was changed every 48hrs in accordance with manufacturer recommendations. Cells were passaged in small clusters using Versene solution and a split ratio of 1:10-1:12. For cryopreservation, cells were suspended in PSC media and temporarily stored in a deep freezer at -80°C before being transferred to liquid nitrogen for long term storage.

***Differentiations and Cell Maintenance:*** 6-well plates were coated with reduced growth factor Cultrex (1mg/12 ml DMEM/F12) at 37°C for at least one hour. PSC were harvested in small clusters using Versene solution without centrifugation, seeded at a passage ratio of 1:12, and cultured in 2 mL of Stemflex medium. After 24 hours, cells were washed with DPBS and 2 mL of appropriate media was added to each differentiating well. For TSC differentiation, TS medium [DMEM/F12 with Glutamax supplemented with 0.1 mM 2-mercaptoethanol, 0.2% FBS, 0.3% BSA, 1% ITS-X supplement, 1.5 µg/ml L-ascorbic acid, 50 ng/ml EGF, 2 µM CHIR99021, 0.5 µM A83-01, 1 µM SB431542, 0.8 mM VPA and 5 µM Y27632] was added to each well. For BMP4 based differentiations 10 ng/ml BMP4 and/or 2uM IWP2 was added to basal differentiation medium (DMEM/F12 with Glutamax supplemented with ITS and L-ascorbic acid 2-phosphate magnesium) was added to each well. Every 24 hours 2 mL of fresh medium was added to each well. Cells were collected on day 2, 4, and 6 for RNA sequencing. For the TS condition, after 6 days of differentiation cells were passaged using TrypLE express at a split ratio of 1:3 and plated on new Cultrex or col IV (5 ug/mL) coated plates. After the first passage, cells were fed every 48 hours and split at a ratio of 1:3-1:6 every 3-4 days. Mature TSC between passages 10-20 were used for all experiments unless otherwise noted. Cells were cultured in humidified incubators 5% $CO_2$ and 20% $O_2$. For cryopreservation, TSC were suspended in Cell Banker 1 and temporarily stored in a deep freezer at -80°C before being transferred to liquid nitrogen for long term storage.

***Single Cell sequencing by Dropseq***

*Dissociation of Cells for Dropseq*

Human iPS cells were grown to 80-90% confluency. After washing cells with DPBS (Thermo Fisher Scientific 14190250) cells were incubated with Versene (Thermo Fisher Scientific 15040066) solution for approximately 10 minutes at 37 °C and gently washed up and down with a P1000 pipette to generate a single cell suspension. Cells were then diluted in Stemflex media and centrifuged at 200g for 4 min before being resuspended in 1 mL of 1X PBS, pH 7.4 (Gibco, # 10010023) with 0.01% BSA (Invitrogen, # AM2616) at the concentration of 100 cells/µL.

*Droplet generation*

Barcoded beads (Chemgenes, # Macosko-2011-10) were washed with 30 mL of ethanol (Sigma Aldrich, # E7023-500ML), then removed all ethanol after centrifugation at 1,000g for 1 minute. Washed beads were suspended in 30 mL of TE-TW (10 mM Tris pH8.0, 1 mM EDTA, 0.5% SDS) and spun down at 1,000g for 1 minute. TE-TW was discarded and re-suspended in 20 mL of TE-TW. 100 µm strainer (Corning, Cat# 352360) was used for size selection of beads.

Beads were suspended as the concentration of 120 beads/µL in 1 mL of the lysis buffer. To generate droplets, cells, beads, and oil (BioRad, #186-4006) were loaded into the 125-µm microfluidic device after aquapel flush treatment (FlowJEM). The cells and barcoded beads suspension were flown at 2 mL/h each and at 13 mL/h for oil.

*Droplet breakage, reverse transcription, and exonuclease I treatment*

Droplets were collected into the 50 mL tubes and 30 mL of 6X SSC was added, then broken with 1 mL of perfluorooctanol (Sigma, # 370533). Collected beads were washed with 1 mL of 6X SSC twice and with 300 µLof RT buffer. On removing the RT buffer, reverse transcription mixture was added into the beads. To synthesize cDNA, beads in RT mix were incubated with rotation at room temperature for 30 minutes and 42 °C for 90 minutes. The reverse transcription mixture was prepared as in the reference, Macosko, E.Z. et. al., Cell 161, 1202-1214 [3]. Beads were washed once with 1 mL of TE-SDS, twice with 1 mL of TE-TW, and with 1 mL of 10 mM Tris pH 8.0. Exonuclease I treatment was performed to remove excess bead primers at 37 °C for 45 minutes with rotation followed by washing beads once with 1 mL of TE-SDS, twice with 1 mL of TE-TW, and with 1 mL of molecular grade water.

*PCR amplification, clean-up with SPRI beads, and library preparation*

Unique barcode STAMPs (Single-cell Transcriptomes Attached to MicroParticles) were resuspended in 1 mL H2O, and loaded on a hemocytometer (Incyto, # DHC-N01). Aliquots of 3,000 beads in 24.6 µL of H2O were amplified with 0.8 µM SMART PCR primer (IDT, 5'-AAG CAG TGG TAT CAA CGC AGA GT-3') and 2X KAPA HiFi Hotstart Ready Mix (Fisher Scientific, # KK2602) using the following PCR condition: (step 1) 95 °C for 3 min, (step 2) four cycles of: 98 °C for 20 sec, 65 °C for 45 sec, 72 °C for 3 min, (step 3) 10 cycles of: 98 °C for 20 sec, 67 °C for 20 s, 72 °C for 3 min, and (step 4) finally, 72 °C for 5 min.

Purified cDNA with 0.6X SPRI beads were analyzed with High Sensitive DNA kit (Agilent, # 5067-4626), then tagmented and amplified using Nextera XT Library Preparation Kit (Illumina, # FC-131-1096) by manufacturer's manual for sequencing.

*Sequencing*

All libraries were checked its quality with Agilent High Sensitive DNA chips before sequencing.

Each library was diluted at 3 nM for an Illumina HiSeq 3000. We used HiSeq SBS 50 cycle kits for sequence (paired-end): 21-bp for read 1 with Custom Read1 primer (IDT, 5'-GCCTGTCCGCGGAAGCAGTGGTATCAACGCAGAGTAC-3'), 21-bp for read 2, and 8-bp for index 1.

*Preprocessing of Drop-seq data*

Raw sequencing data was preprocessed using the pipeline "Drop-seq Alignment Cookbook" v2.0.0 found at https://github.com/broadinstitute/Drop-seq/releases/ and described in [3]. Briefly, paired-end reads were filtered in order to remove read pairs with any base with quality of less than 10 in both cellular and molecular barcodes. SMART adapters at 5' end and polyA tails at 3' end with 6 or more bp were removed from the second pair, and then aligned to the reference human (GRCh38) genome using HISAT2 v2.1.0 [4] with the default settings. Uniquely mapped reads were retained and grouped by cell barcode. UMI barcodes with a hamming distance = 1 were merged within each cell, and the number of transcripts of each gene was defined based on the total number of each distinct UMI sequence, for that gene, in a given cell.

*Clustering and Differential Gene Expression analysis for Marker Genes*

All downstream analysis for quality control, clustering, and differential expression were performed using Seurat [5] v3.0 R package. Data was initially filtered to remove low quality cells and very low detected genes. We discarded cells with less than 1000 genes detected and with more than 20% of mitochondrial gene content. Genes expressed in less than 3 cells were also removed. Variance stabilizing and normalization was performed using sctransform [6] approach adjusting for library size and the percentage of mitochondrial genes in each cell. To cluster the cells, cell distances were defined with 30 first PCs using a graph-based approach implemented in Seurat, that briefly build a K-nearest neighbor (KNN) graph based on euclidean distance in PCA space, with subsequent refining of edge weights using Jacard similarity. Clusters were defined using Seurat default algorithm – a shared nearest neighbor (SNN) graph based on k-nearest neighbors with modularity optimization using original Louvain algorithm. Gene markers of cell clusters were defined using ROC analysis for only genes with at least 0.25 logFC average difference between the clusters. Graphic visualizations of clusters and gene expression were done using Seurat and ggplot2 in R.

*Single-cell trajectory reconstruction*

Reconstruction of cell differentiation trajectories for cell clusters was performed with STREAM (Single cell Trajectories Reconstruction, Exploration and Mapping) [7] Python package. The expression data of high-quality cells (9,821 cells) was library size normalized, scaled, natural-log transformed and used as input for STREAM. Data dimension reduction was done with Spectral Embedding (SE) based on most variable genes found with LOESS fitting. Branches representing trajectory structure were learned based on 3 SE dimensions.

Marker genes of each branch were defined by Leaf gene detection analysis implemented in the STREAM package. Initially, the average expression of each gene is calculated for all branches. Based on the average expressions, Z-scores are computed and genes are picked as candidates when greater than 1.5. Then a Kruskal–Wallis H-test is calculated for each candidate to test the difference of gene expression among branches. Finally, the Conover test for multiple comparison is computed for the differentially expressed candidates (Krustal-Wallis p-value < 0.01) between all branches, and genes with Conover p-value < 0.01 are picked as a Leaf gene for that branch. GO enrichment analysis of branch markers were performed for top 100 leaf genes using hypergeometric test with GOstats v2.48.0 R package. Circus plot with GO terms was done using GOplot v1.0.2 R package.

Transition genes were found by performing transition gene analysis implemented in STREAM. The algorithm starts ordering cells based on their inferred pseudotime and splits them in two groups. One group contains 20% of cells closer to the pseudotime start point, and the other is composed of all remaining cells (80%). Gene average expressions are calculated for each group and genes with a log2 fold change greater than 0.25 between groups are picked as candidate genes. Then, Spearman's rank correlation is computed between inferred pseudotime and gene expression of the candidates. Genes with correlation coefficient above 0.4 are reported as transition genes.

Overrepresentation analysis was performed by using the algorithm MSET [8] implemented in R, using 10000 permutations. The list of transcription factors related to trophoblast differentiation used is described in [9].

*Enrichment analysis for placental cell-specific genes.*

To characterize the expression profiles of the 19 clusters detected in this work, we analyzed the overlap with cell-specific genes identified by previous single-cell studies. We used two sets of candidate genes. A set of 38 cell-specific gene lists (30 gene each) was drawn from the supplementary table 2 of [10]. These gene lists span both maternal (decidua and blood) and fetal tissues (placenta). Another set of 14 cell-specific gene lists was drawn from the supplementary table 1 of [11], which also span analogous tissues.

To profile the expression patterns of the 19 clusters detected in this work, we applied the Cell-Specific Expression Analysis (CSEA) as implemented in the pSI R package [12]. This approach compares each profile with all the others and identifies genes expressed in one group but not in the others, calculates a score (specificity index, SI) for each gene and attributes a statistical level of significance (pSI). Here, we used the normalized expression matrix to compute cluster-level median gene expression for each gene, which defined 19 cluster profiles. Then, cluster profiles were used to compute SI and pSI for each gene, within each profile. Cluster-specific gene lists were obtained applying the statistical threshold, pSI < 0.05.

Finally, we tested whether cell-specific genes from previous studies are over-represented in our clusters. We used the hypergeometric test and applied the Bonferroni correction for multiple comparisons, considering all the tested gene lists [$\alpha = 0.05/ (19\times (38+14)) = 5.1\times10^{-5}$].

In order to compute the similarity of the cells for different developmental stages we used gene lists defined as cell type-specific by Xiang et al. [13]. Counts matrix was filtered to keep the same cells and genes used in clustering analysis and the data was normalized using the function NormalizeData from Seurat v.4.0.2 with scale factor equal to 10,000. The similarity to each gene expression program was computed using the function AddModuleScore from Seurat. Basically, the function calculates the average expression levels of each group of cell type-specific genes (programs) on a single cell level, then subtract by the aggregated expression of control feature sets.

Comparisons of gene expression levels for cell type-specific genes among clusters were done by using average expression of each gene and performing pairwise Wilcoxon Rank Sum test. P-values were combined by using the Holm-middle method implemented in the function combinePValues from scran v.1.18.7 R package.

Differential expression analysis for amnion genes was performed by the function FindAllMarkers from Seurat R package using Wilcoxon Rank Sum test with Bonferroni correction.

*PANDA regulatory networks*

Gene regulatory networks were constructed using the pandaR v1.14.0 R package which integrates multiple types of data to infer direct interactions (edges) between TFs and target genes. PANDA initiates with a prior regulatory network that can be built by mapping TF binding sites to the genome, and refines this initial network integrating gene expression data of target genes and TF PPI data. The main idea behind the algorithm is that target genes from a given TF are likely co-expressed, and TFs that interact with each other are more likely to regulate a similar group of target genes. These two assumptions are used to infer the edge weight of each TF-target interaction, and basically reflects the congruence between the regulatory profile of a TF with target gene co-expression. Iteratively the algorithm refine the initial network structure and infer a final consensus regulatory network.

A regulatory network was built for each cluster using as initial network a TF-motif binding map described in [14] and downloaded in (https://sites.google.com/a/channing.harvard.edu/kimberlyglass/tools/resources). Gene expression data was library size normalized, scaled, and natural-log transformed. Genes expressed in less than 10% of the cells of each cluster were filtered out. PPI network of TFs was built using STRING database v11.0 (downloaded from https://string-db.org/) and score interactions were divided by 1000 to initiate the PANDA.

Edges with negative weights estimated by PANDA were discarded in order to explore only interactions with greater evidence. The relative importance of individual genes in the network topology was addressed with the betweenness centrality measure calculated using the Igraph v1.2.4 R package. For each cluster, the significance of betweenness values were calculated based on the betweenness probability distribution of 100 random networks constructed with the same number of edges of that cluster. Graphic visualization, gene clusterization, and GO enrichment analysis of gene clusters within networks were performed using Cytoscape v3.7.2 [15] and ClueGO v2.5.4.ENRICHMENT/RANKING ANALYSIS [16].

*Software and Analysis:*

Analysis pipelines are publicly deposited on Github:https://github.com/paquolalab/placenta_ips

*EVT and STB differentiation*

Differentiation of TSCs were performed as previously described [17], with minor modifications. For EVT differentiation, 24-well plates ibidi were coated with 1 µg/mL Collagen IV overnight. $2 \times 10^5$ TSC were seeded per well in 500µL EVT basal medium [DMEM/F12 supplemented with 0.1 mM β-mercaptoethanol, 0.5% penicillin-streptomycin, 0.3% BSA, 1% ITS-X, 7.5 µM A83-01, 2.5 µM Y27632] supplemented with 4% KSR and 100 ng/mL NRG1 alpha. Matrigel was added to a 2% final concentration shortly after resuspending TSC in the medium. On day 3, the media were replaced with 500µL EVT basal medium supplemented with 4% KSR, and Matrigel was added to a 0.5% final concentration. At day 6, the media were replaced with 500µL EVT basal medium, and Matrigel was added to a 0.5% final concentration. At day 9, the cells were ready for analysis

For STB differentiation, 24-well plates ibidi were coated with 2.5 µg/mL Collagen IV overnight. $2 \times 10^5$ TSCs were seeded per well in 500µl STB medium [DMEM/F12 supplemented with 0.1 mM β-mercaptoethanol, 0.5% penicillin-streptomycin, 0.3% BSA, 1% ITS-X, 2.5 µM Y-27632, 2 µM Forskolin and 4% KSR]. The media was changed at day 3, and at day 4 the cells were ready for analysis.

*Flow Cytometry*

Trophoblast stem cells-like were dissociated with TrypLE, passed through a 70 mm mesh filter, and suspended in 2% BSA/PBS. For flow cytometric analysis of ITGA6, cells were incubated with FITC-conjugated anti-CD49f (Integrin alpha 6 - ITGA6) for 30 minutes at room temperature and then fixed with 4% PFA for 15min. FITC-conjugated rat IgG2a was used as isotype control. Flow cytometry was carried out using the Becton Dickinson LSRII and the acquisition was done in BD FACSDiva Software Version 8.0.1. The data were analyzed using FlowJo software.

*Immunofluorescence Staining*

Cells were washed in Tris Buffered Saline (TBS), fixed with 4% paraformaldehyde for ten minutes, and washed three times in TBS. Cells were then incubated in blocking solution (3% horse serum, 0.1% Triton-X in TBS) for one hour at room temperature before being incubated with primary antibodies in blocking solution overnight at 4°C. Cells were then rinsed twice and incubated with secondary antibodies in blocking solution for two hours at room temperature. After two washes in TBS cells were counterstained with Hoechst 33342 and rinsed twice in TBS. Cells were imaged using a Zeiss LSM770 or LSM780 confocal microscope. The brightness and/or contrast was globally adjusted for some images.

*HLA-G and Fusion Index Analysis*

The HLA-G positive cells were calculated using the following formula: [(EVT-N)/T x 100]). EVT, the number of HLA-G positive cells; N, the number of HLA-G negative cells; T, total number of nuclei counted. The fusion index represents the percentage of cell-cell fusion events. Syncytia were defined as cells with at least three nuclei. The number of Hoechst 33342I-stained nuclei and syncytia were counted using Image-J. The fusion index was calculated using the following formula: [(N-S)/T] x 100. N, the number of nuclei in the syncytia; S, the number of syncytia; T, total number of nuclei counted.
